# Mammary Fibroblasts Secrete Damage Associated Molecular Patterns through Extracellular Vesicles in Response to Ionizing Radiation

**DOI:** 10.1101/2025.11.25.690532

**Authors:** Greg Berumen Sánchez, Purvi Patel, Preston Gomez-Crase, Chloe Kim, Kristie Lindsey Rose, Marjan Rafat

## Abstract

Ionizing radiation (IR) is an integral component of cancer therapy. Cellular exposure to IR typically leads to major biological consequences including cell death and senescence. Furthermore, tissue injury in known to involve the release of damage-associated molecular patterns (DAMPs) into the extracellular space, which trigger inflammation and wound healing. However, DAMP release in the context of radiation injury remains to be fully characterized. Evidence suggests that extracellular vesicle (EV) secretion and associated cargo components are part of the cellular response to IR, but the mechanisms integrating cellular damage and EV secretion post-IR are largely unexplored. In this study, we show that acute IR-induced damage in mammary fibroblasts results in a senescence-like phenotype and substantially increased EV secretion. Quantitative proteomic analysis revealed that IR-induced EVs are enriched with extracellular and intracellular DAMPs, along with other pro-inflammatory mediators. We show that knockdown of the GTPase Rab27a abrogates IR-induced EV secretion and inhibits the enrichment of key DAMPs in EVs. By examining the integration of cellular damage and senescence with the release of inflammatory signals, this study elucidates a potentially critical role for EV-associated proteins in the radiation response.

## 1 INTRODUCTION

Cancer is the leading cause of premature death in 57 countries, including the United States, China, and much of Europe (Bray et al., 2021), and it may soon surpass cardiovascular disease as the leading cause of premature death in most countries (Bennett et al., 2020). In 2022, approximately 19 million new cancer cases were diagnosed, with about 10 million cancer-related deaths (Bray et al., 2024). Radiotherapy is a predominant cancer treatment modality and it is estimated that over 50% of all cancer patients benefit from radiation for palliative and curative care (Yap et al., 2016). In breast cancer patients, radiotherapy has an 87% utilization rate, the highest among the most frequent cancers (Atun et al., 2015; Laskar et al., 2022).

Previous clinical investigations from our group focusing on breast cancer patients have uncovered a connection between radiotherapy and cancer recurrence, particularly in immunocompromised patients (Rafat et al., 2018; Sherry et al., 2020). These clinical findings were corroborated by *in vivo* studies, where radiation of normal mammary tissue resulted in macrophage infiltration and circulating tumor cell (CTC) recruitment (Hacker et al., 2023; Rafat et al., 2018). However, further examination of the irradiated breast stroma is needed to improve our understanding of the key cellular and molecular processes governing alterations in intercellular communication and how that communication dictates tissue development and patient outcome.

Within the tissue microenvironment, intercellular communication is conducted through juxtracrine interactions, where cells exchange signals through direct contact, or by paracrine interactions, where cells communicate by secreting signals into the extracellular space (Gilbert, 2003). Vesicle-mediated communication through the secretion, interaction, and uptake of membrane-bound particles broadly termed extracellular vesicles (EVs), has emerged as a primary focus of the field of intercellular communication (van Niel et al., 2022; Wessler & Meisner-Kober, 2025). EVs are lipid-bilayer delimited nanoparticles that are secreted by all cell types and organisms and have been shown to contain all major classes of biological molecules. EVs can interact with target cells and transmit biologically active cargo, including proteins, nucleic acids, lipids, and metabolites (Colombo et al., 2014; C. Williams et al., 2019). The lipid-bilayer membrane protects key cargo components from degradation, allowing for signaling factors to survive the harsh conditions of the extracellular space and avoid unintended immune surveillance (Pua et al., 2019). Additionally, EVs are understood to possess cell-specific targeting capabilities, further highlighting their ability to serve as primary drivers of crosstalk between cells in the local microenvironment and in distal sites (Choi et al., 2024). While EV-mediated communication is imperative for normal physiology and homeostasis, alterations in EV secretion and cargo components are common in the context of tissue injury and disease (Berumen Sánchez et al., 2021).

Breast cancer predominantly arises in the terminal duct lobular units, or lobules, within breast tissue (Russo et al., 2000). Histologically, lobules contain a fibroblast rich stroma (Sumbal et al., 2021), and clinical analysis of breast tissue specimens reveals fibrosis, atrophy, and atypical fibroblasts as consistent radiation-induced pathological changes (Moore et al., 2004; Schnitt et al., 1984; Straub et al., 2015). Therefore, investigating the contribution of fibroblasts to the alterations within the breast tissue microenvironment is crucial in understanding how the molecular effects of ionizing radiation (IR) may impact patient outcomes. In this work, we investigate the cellular and molecular effects of IR on intercellular communication within the breast stroma. We focus on the IR-induced secretion and characterization of EVs from mammary fibroblasts and their implications in the intercellular communication landscape.

## 2 MATERIALS AND METHODS

### 2.1 Cell culture

Human mammary fibroblasts originally isolated from reduction mammoplasty tissue (Kuperwasser et al., 2004) were obtained in May 2022 and maintained in high glucose DMEM (#11995065, Gibco) containing 10% bovine calf serum (Gibco) and 1X penicillin-streptomycin (P/S; Gibco). Cells were grown on tissue culture-treated flasks in a humidified, 5% CO_2_ incubator at 37°C. Culture medium was replaced every 2–3 days. Cell viability was assessed via trypan blue exclusion. Briefly, equal parts of cell suspension and 0.4% trypan blue (Invitrogen) were combined, and the concentration of total and live cells was measured using a TC-20^TM^ cell counter (Bio-Rad). A lentiviral short hairpin RNA (shRNA) expression system, pLKO.1 (Moffat et al., 2006), was used to knock down p53 [TRCN0000003753 (5’-CCGG-CGGCGCACAGAGGAAGAGAAT-CTCGAG-ATTCTCTTCCTCTGTGCGCCG-TTTTT-3’), Sigma Aldrich], Rab27a [TRCN0000005296 (5’-CCGG-CGGATCAGTTAAGTGAAGAAA-CTCGAG-TTTCTTCACTTAACTGATCCG-TTTTT-3), Sigma Aldrich], or a non-targeting scrambled control [#SHC002 (5’-CCGGCAACAAGATGAAGAGCACCAACTC-GAGTTGGTGCTCTTCATCTTGTTGTTTTT-3’), Sigma Aldrich]. Lentiviral particles were generated using HEK-293T cells via co-transfection with shRNA and packaging plasmids (psPAX2, pMD2.G) following standard protocols (Addgene, 2006). Briefly, lentiviral particle-containing conditioned media (CM) was filtered using a 0.45 μm PES syringe filter and kept at 4°C for 1–2 days or stored at −80°C. 1 mL lentiviral particles was used to infect target cells in 6 cm dishes for 24 hours. Cells were allowed to rest for an additional 24 hours post-infection by removing CM and replacing with fresh growth medium. After the 24 hour rest period, CM was replaced with complete growth medium containing 4 μg/mL puromycin (Sigma) for selection of transformed cells. CM was replaced with fresh puromycin-containing medium every other day. After 3–5 days of selection, transformed cells were analyzed via western blot and/or cultured in complete growth medium containing 1 μg/mL puromycin to stably maintain knock downs.

### 2.2 Radiation

Cells were cultured for 2–3 days prior to treatment with γ-radiation using a Mark I Model 68 Cesium-137 Irradiator (JL Shepherd and Associates). Cells were irradiated to a single dose of 10 Gy on a spinning platform at a rate of 1.86 Gy / min, and controls were sham treated. Cells were rinsed 2x with calcium- and magnesium-free Dulbecco’s PBS (Gibco) prior to replacing medium with serum-free Opti-MEM (#31985070, Gibco) containing 1X P/S. Cells were placed in a humidified, 5% CO_2_ incubator at 37°C. Culture medium was not changed after replacing with Opti-MEM.

### 2.3 Extracellular vesicle (EV) isolation

Cell cultures at 60–80% confluence were incubated in serum-free Opti-MEM for 48 hours. For each replicate and treatment condition, 200 mL of CM were collected from approximately 2–4×10^7^ cells. Cell counts were determined following trypsinization (Gibco) to estimate EV secretion rates, calculated as particle concentration per cell per hour. CM was subjected to serial centrifugation at 300×*g* for 5 min, 2,000×*g* for 20 min, 10,000×*g* for 37 min and 100,000×*g* overnight to sediment live cells, dead cells and debris, large EVs, and small EVs, respectively (**Figure 1a**). All centrifugation steps were performed at 4°C with maximum acceleration and deceleration settings. Centrifugation steps through 10,000×*g* were performed in a Sorvall® ST40R (Thermo Scientific) centrifuge. The 300×*g* and 2,000×*g* steps were performed using a TX-1000 (K-Factor=9,469) swinging bucket rotor (Thermo Scientific) and 50 mL conical polypropylene tubes, approximately 40 mL CM/supernatant per tube. The 10,000*g* step was performed using a Fiberlite^TM^ F15-6×100y (K-Factor=2,045) fixed-angle rotor (Thermo Scientific) and Oak Ridge polycarbonate tubes (#3118-0085, Thermo Scientific), approximately 66 mL supernatant per tube. Ultracentrifugation was performed using an Optima^TM^ L-90 or Optima XPN floor-model ultracentrifuge (Beckman Coulter) and Type 45 Ti (K-Factor=133) fixed-angle rotor (Beckman Coulter) with polycarbonate bottle assemblies (#355622, Beckman Coulter), approximately 66 mL supernatant per tube. For each sample, the pellets from 3 tubes were combined by re-suspending in 3 mL ice-cold PBS. Re-suspended pellets were spun again at 100,000×*g* for 1–3 hours in an Optima TL-100 or Optima MAX-TL (Beckman Coulter) table-top ultracentrifuge using TLA100.3 (K-Factor=12) fixed-angle rotor (Beckman Coulter) and thickwall polycarbonate tubes (#349622, Beckman Coulter). Washed EV pellets were resuspended in 100 μL PBS containing 5 mM EDTA for downstream particle analysis, or 20–100 μL lysis buffer (5% SDS, 10 mM EDTA, 120 mM Tris-HCl, pH 8.0) and incubated at 200 RPM on an orbital shaker for ≥ 15 min (up to overnight) for protein extraction. PBS samples were used immediately or kept at 4°C for ≤ 7 days. Concentration of protein extracts was determined through bicinchoninic acid assay (Thermo Scientific), and samples were stored at −80°C or used immediately.

**Figure 1.**
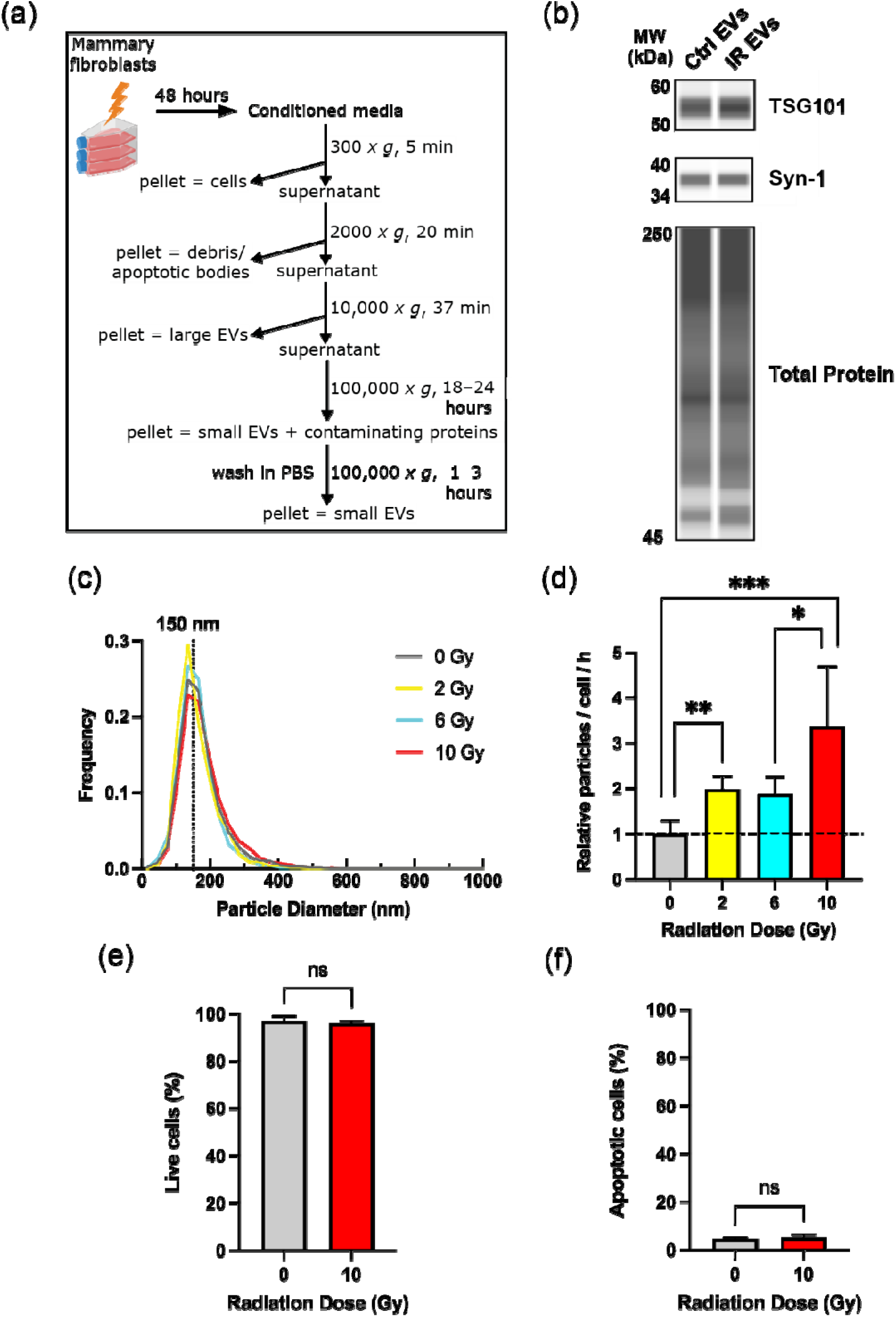
γ**-radiation induces secretion of EVs but not cell death or apoptosis in RMFs. (a)** Schematic of the experimental procedure for isolation of EVs from irradiated RMF cultures. **(b)** Representative western blot analysis of syntenin-1 (Syn-1) and tumor susceptibility gene 101 (TSG101) expression in EV-derived protein extracts from control (Ctrl; 0 Gy) and irradiated (IR; 10 Gy) RMFs. **(c)** Size distribution of EVs from RMFs treated with the indicated IR dose (N=3), obtained via nanoparticle tracking analysis (NTA). Approximate peak diameter is indicated with dotted line. **(d)** Particle concentration normalized to cell number per hour for EVs isolated from RMFs treated with the indicated IR dose (N=3). Data is relative to control average. Statistical analysis was performed using unpaired *t*-tests with Welch’s correction. **(e)** Percentage of live cells in RMF cultures 48 hours post-IR, determined through trypan-blue exclusion of dead cells (N=3). Statistical analysis was performed using Mann-Whitney test. **(f)** Percentage of apoptotic (Annexin V+) RMFs 48 hours post-IR, determined via flow cytometry (N=3). Statistical analysis was performed using Mann-Whitney test. Statistical significance was determined as described with nonsignificant (ns) *p* > 0.05, \**p* < 0.05, \*\**p* < 0.01, and \*\*\**p* < 0.001. Error bars show standard deviation.

### 2.4 Nanoparticle tracking analysis (NTA)

NTA was performed using the ZetaView® PMX-120 (Particle Metrix) equipped with a 488 nm laser. In each case, 1 mL of sample diluted in 0.2 μm-filtered PBS was measured by the instrument. Typical dilutions were between 1:1,000–1:10,000. In some experiments, samples were treated with Triton® X-100 (Sigma Aldrich) to a final concentration of 0.1% for 10 min at room temperature prior to dilution in PBS and measurement. Particle size distribution and sample concentration were determined through the instrument software (v8.05.12 SP2). For each measurement, the instrument was set to sensitivity=80, frame rate=30 and shutter speed=70. Post-acquisition parameters were set to minimum brightness=20, maximum size=1,000 pixels, minimum size=5 pixels, and tracelength=15. Polystyrene beads (#23A1009, Alpha Nano Tech) with average size of 100 nm were used to calibrate the instrument prior to each experiment.

### 2.5 Apoptosis detection via flow cytometry

Cell cultures were digested into single cell suspensions via trypsinization, rinsed with DPBS, and re-suspended at a concentration of 1–10×10^6^ cells/mL in annexin binding buffer (10 mM HEPES, 140 mM NaCl, and 2.5 mM CaCl_2_, pH 7.4). For each sample, 100 μL of cell suspension was aliquoted into a 96-well round bottom microplate (Falcon) and incubated with 5 μL Alexa Fluor 647-conjugated Annexin V (A23204, Invitrogen) for 15 min at room temperature, protected from light. After staining, cells were immediately analyzed using a 4-laser Amnis® CellStream® flow cytometer (Cytek Biosciences). Data analysis was performed using FlowJo^TM^ v10 software.

### 2.6 Negative-stain transmission electron microscopy (TEM)

5 μL of EV samples were adhered to freshly glow discharged carbon coated 300 mesh Cu grids (Electron Microscopy Sciences) for 20 sec followed by negative staining using 2% uranyl acetate (Electron Microscopy Sciences). TEM was conducted using a FEI Tecnai G2 Osiris system.

### 2.7 Western blot

Immunoassay analysis was primarily conducted using the Jess^TM^ Western Blot System (ProteinSimple) using the 25-capillary cartridge and 12–230 kDa separation module (SM-W004) according to the manufacturer’s protocol. Briefly, protein extracts were diluted with 0.1X sample buffer and combined 4:1 with 5X fluorescent master mix containing 200 mM dithiothreitol, to a final protein concentration of 0.05–0.2 mg/mL. Samples were vortexed and heated at 95°C for 5 min, then loaded into the assay plate along with the molecular weight ladder, blocking reagents, primary antibodies, and HRP-conjugated secondary antibodies (ProteinSimple). RePlex^TM^ (RP-001) and total protein detection modules were utilized (DM-TP01) for data normalization. Electrophoresis, blocking, antibody incubations, washes, and immuno- and total protein detection were conducted through the Jess system with default settings. Data were analyzed using Compass v6.1.0 (ProteinSimple). Default parameters were used for peak fit analysis, with ‘Dropped Lines’ selected for peak area calculation. Normalization was enabled and region was set to Area Start=0.0 kDa, Area End=250.0 kDa. Target peak area and total protein data are presented as pseudo-blots in the manuscript. Primary antibodies used: Anti-Syntenin (ab133267, Abcam; 1:20), anti-TSG101 (ab30871, Abcam; 1:20), anti-p21 (#2947, Cell Signaling; 1:50), anti-p53 (MA5-12557, Invitrogen; 1:20), anti-GAPDH (#5174S, Cell Signaling, 1:20), anti-β actin (AM4302, Invitrogen, 1:50), anti-SERPINE2 (#11303-1-AP, Proteintech, 1:20).

Traditional immunoblots were conducted when deemed unsuitable for the Jess system. 5–20 μg of protein extract was diluted in RIPA buffer (Thermo Scientific) and combined 3:1 with 4X sample loading buffer (LI-COR Biosciences) containing 355 mM 2-mercaptoethanol (Bio-Rad). Samples were vortexed and heated at 95°C for 5 min. Samples were then loaded onto a 4–20% tris/glycine polyacrylamide gel (Bio-Rad) along with pre-stained protein standards (Bio-Rad). Protein separation was achieved using tris/glycine/SDS buffer system by supplying 250 V for 25–30 minutes at room temperature. Protein was transferred to a 0.45 μm pore size PVDF membrane (Bio-Rad) via tank blotting method using Towbin buffer by supplying 30 V for 16 hours at 4°C. Following transfer, membranes were dried for 10 min at 37°C then incubated in methanol for 30 sec, washed with TBS for 5 min, and stained with Revert^TM^ 700 (LI-COR Biosciences) Total Protein Stain according to the manufacturer’s protocol. Membranes were imaged in the 700 nm channel of the Odyssey FC Imaging System (LI-COR Biosciences) with an exposure time of 30 sec. Membranes were subsequently blocked for 1 hour at room temperature using Intercept® TBS Blocking Buffer (LI-COR Biosciences) followed by primary antibody incubation overnight at 4°C. Membranes were washed 4x, 5 min each, using TBS containing 0.2% Tween^TM^ 20 (Invitrogen) and then incubated with an HRP-conjugated secondary antibody (Promega) at 1:2,500, IRDye® 800- and/or IRDye 680-conjugated secondary antibodies (LI-COR Biosciences) at 1:15,000. Membranes were imaged in the chemiluminescence and 600, 700, and 800 nm channels with 30 sec or 2 min exposure time, as necessary. Data were analyzed using Image Studio v6.0 (LI-COR Biosciences). Analysis type was ‘Manual’ with ‘User-Defined’ background subtraction method. Primary antibodies used: anti-CD63 (ab134045, Abcam; 1:1,000), anti-GM130 (NBP2-53420, Novus; 1:1,000), anti-Rab27a (#69295S, 1:1,000), anti-β actin (AM4302, Invitrogen, 1:1000), anti-Histone H4 (#39269, Active Motif, 1:1,000), anti-LAMB2 (#30943-1-AP, Proteintech; 1:5000), anti-LC3B (ab192890, Abcam, 1:1000).

### 2.8 LC–MS/MS and data analysis

EV-derived protein extracts were reduced with 10 mM TCEP (tris(2-carboxyethyl)phosphine) at 55°C for 15 min and alkylated with 20 mM iodoacetamide for 30 min at room temperature, protected from light. Protein samples were then prepared by S-Trap (ProtiFi) digestion. Aqueous phosphoric acid was added to each sample to a final concentration of 2.5%, followed by addition of S-Trap binding/wash buffer (90% methanol containing 100 mM tetraethylammonium bromide (TEAB)) at 6X the sample volume. Samples were loaded onto S-Trap micro columns and centrifuged at 4,000×*g* followed by 4 washes of 150 µl each using binding/wash buffer. Proteins were digested with trypsin (Promega; 1:10 enzyme to protein ratio) in 50 mM TEAB, pH 8.0, for 1h at 47°C. Peptides were eluted by sequential addition of 40 µL each of 50 mM TEAB, 0.2% formic acid, and 0.2% formic acid in 50% acetonitrile and were dried in a speed-vac concentrator. Peptides were next analyzed using a tandem mass tag (TMT) approach. TMT-based quantitative proteomics analysis was performed using TMTsixplex™ Isobaric Label Reagent Set (#90066, Thermo Scientific). Peptides were reconstituted in 50 mM TEAB and labeled with TMT tags following the manufacturer’s instructions. Labeled peptides (7 μg per sample) were combined and fractionated using the Pierce High pH Reversed-Phase Peptide Fractionation Kit (#84868, Thermo Scientific) according to the manufacturer’s protocol for TMT-labeled peptides. Elution steps (8 fractions) consisted of the following: 10%, 12.5%, 15%, 17.5%, 20%, 22.5%, 50%, and 60% acetonitrile with 0.1% triethylamine. The eluted fractions were dried and reconstituted in 0.2% formic acid for LC–MS/MS analysis. A reverse phase capillary column was packed with 20 cm of C18 material (Jupiter, 3 μm beads, Phenomenex) as described previously (Howard et al., 2024). Peptides were loaded on column using a Dionex Ultimate 3000 nanoLC and autosampler. Mobile phase solvents consisted of 0.1% formic acid, 99.9% water (solvent A) and 0.1% formic acid, 99.9% acetonitrile (solvent B). Peptides were gradient eluted at a flow rate of 350 nL/min and were analyzed using varied reverse phase gradients over 90 min. For fraction 1, peptides were analyzed with the following gradient: 5–18% B in 75 min, 18–50% B in 6 min, 50–70% B in 3 min, 70–2% B in 1 min, 2% B for 5 min (column re-equilibration). For fractions 2–5, the first 2 steps of the gradient were adjusted to 5–25% B in 75 min and 25–50% B in 6 min, and the following 3 steps were identical to fraction 1. For fraction 6, the gradient included 5–30% B in 75 min, 30–50% B in 6 min, and the following 3 steps were identical to fractions 1–5. For fractions 7 and 8, the gradient was as follows: 10–50% B in 75 min, 50–90% B in 8 min, 90% B for 1 min, 90–2% B in 1 min, 2% B for 5 min. Peptides were analyzed using a data-dependent acquisition method on an Orbitrap Exploris 480 mass spectrometer (Thermo Scientific), equipped with a nanoelectrospray ionization source. The instrument method consisted of MS1, followed by up to 20 MS/MS scans, with an AGC (automatic gain control) target of 2×10^5^. Higher-energy collisional dissociation (HCD) was set to 35 nce, and dynamic exclusion (15 sec) was enabled.

Data were searched in Proteome Discoverer 2.2 (Thermo Scientific) using SequestHT for database searching against a UniProtKB human protein database. Default workflow templates in Proteome Discoverer were applied and included the “PWF QE Reporter Based Quan SequestHT Percolator” processing and the “CWF Comprehensive Enhanced Annotation Reporter Quan” consensus workflows. Search parameters included trypsin cleavage with two missed cleavage sites, carbamidomethyl (C) and TMTsixplex (K, N-terminus) as static modifications, and dynamic modification of oxidation (M) and acetylation (N-terminus). Percolator validation was performed with a target FDR setting of 0.01 for high confidence identifications. During quantitative analysis in the consensus workflow, data were normalized and summed abundance-based ratios were used for ratio calculations. TMT results were also filtered to include proteins with a minimum of two unique peptides. The background-based ANOVA method was selected for statistical analysis. Quantified proteins with absolute log_2_ abundance ratio ≥ 0.59 and adjusted *p-*value < 0.05, were chosen to be significant. Significant proteins with %CV greater than the upper outlier threshold, as determined via the Generalized Extreme Studentized Deviate test (MATLAB_R2024a), were excluded.

### 2.9 Detection of γH_2_AX foci

Cells grown on coverslips were irradiated or sham treated then incubated in a humidified, 5% CO_2_ incubator at 37°C for 30 min. Cells were fixed for 15 min with 10% neutral buffered formalin (VWR) and permeabilized for 10 min with 0.1% Triton X-100 in PBS. Coverslips were washed 2x with PBS and incubated for 1 hour at room temperature in blocking buffer (1% BSA (Sigma Aldrich), 22.52 mg/mL glycine (RPI) in PBS-T (PBS containing 0.1% Tween 20)). Then, cells were incubated with anti-phospho-Histone H2A.X (MA5-38225, Invitrogen) at 1:100 in blocking buffer overnight at 4°C in a humidified chamber. Following overnight incubation, cells were washed 2x for 5 min with PBS-T and incubated with an anti-rabbit IgG Alexa Fluor 594 conjugated secondary antibody (A-21207, Invitrogen) at 1:200 in blocking buffer for 1 hour at room temperature, protected from light. Coverslips were washed 2x for 5 min with PBS-T and mounted on microscope slides (VWR) using NucBlue^TM^-containing mounting media (Invitrogen) and let to dry overnight, protected from light. Images were acquired with a DMi8 inverted microscope (Leica) at 100X magnification using 63x/1.40 NA Plan Apo Oil objective lens (Leica) and DFC3000 G camera (Leica).

### 2.10 Senescence associated β-Galactosidase (SA-βgal) activity assay

SA-βGal activity was measured using the Senescence β-Galactosidase Staining Kit (Cell Signaling) following the manufacturer’s instructions. Briefly, cells were fixed at room temperature for 10 min with the supplied fixative and incubated for 24 hours at 37°C with X-gal staining solution (pH 6.0). Brightfield images were obtained on a DMi8 inverted microscope (Leica) at 10X magnification using HC PL Fluotar 10x/0.32 dry objective (Leica) and MC190 HD camera (Leica).

### 2.11 5-Ethynyl-2’-deoxyuridine (EdU) incorporation assay

EdU incorporation was detected using the Click-iT® Plus EdU Imaging Kit (Invitrogen) following the manufacturer’s instructions. Briefly, cells cultured on chambered coverglass (Millicell® EZ Slide, Millipore) were incubated in growth medium containing 10 μM EdU in a humidified, 5% CO_2_ incubator at 37°C for 1 hour. Following incubation, cells were fixed in 10% neutral buffered formalin (VWR) for 15 min at room temperature. Cells were washed 2x with PBS and permeabilized with 0.5% Triton X-100 in PBS for 20 min at room temperature. Then, cells were washed 2x in PBS and incubated with the supplied reaction cocktail for 30 min at room temperature, protected from light. Reaction cocktail was removed, and cells were washed with PBS. Media chamber was removed from coverglass and a coverslip was mounted using NucBlue-containing mounting media (Invitrogen) and let to dry overnight, protected from light. Images were acquired with a DMi8 inverted microscope (Leica) at 100X magnification with 63x/1.40 NA Plan Apo oil objective lens (Zeiss) and DFC3000 camera (Leica).

### 2.12 Luminex cytokine array

Specimens for analyte testing were collected from control and irradiated cell cultures and filtered using a 0.45 μm PES syringe filter. For each replicate and experimental condition, 200 μL of filtered CM was submitted to Eve Technologies Corp. (Calgary, AB, Canada). Analysis of specimens was performed using commercially available MilliporeSigma MILLIPLEX^®^ base kits, using the Luminex 200™ system (Luminex) with Bio-Plex Manager™ (BPM) software (BioRad). Fifteen markers were simultaneously measured in the samples using Eve Technologies’ Human Focused 15-Plex Discovery Assay®, according to the manufacturer’s protocol. Markers for the 15-plex assay are listed in **Table S1**. Assay sensitivities of these markers range from 0.14–5.39 pg/mL. Individual analyte sensitivity values are available in the MilliporeSigma MILLIPLEX® MAP protocol. Thirteen markers were simultaneously measured in the samples using Eve Technologies’ Human MMP/TIMP 13-Plex Discovery Assay® which consists of two separate kits: one 9-plex and one 4-plex (R&D Systems), both utilized according to the manufacturer’s protocol. Combined markers for the 13-plex assay are listed in **Table S2**. Assay sensitivities of these markers range from 0.28–253 pg/mL. Individual analyte sensitivity values are available in the product datasheet for the 4-Plex and by building the panel on the R&D Systems Magnetic Luminex Performance product page for the 9-Plex.

### 2.13 p53 activation

Cells were treated with 10 μM Nutlin-3a (Sigma) in complete growth medium for 72 hours. Control cells were treated with an equal volume of DMSO. After treatment, culture medium was replaced with OptiMEM for EV isolation as described in **2.3**, or cells were rinsed with ice-cold PBS and whole cell protein extract was collected in RIPA buffer.

### 2.14 Macroautophagy flux analysis

Macroautophagic flux was determined via Western blot analysis of microtubule-associated proteins 1A/1B light chain 3B (LC3) in protein extracts derived from cells treated with or without an inhibitor of autophagy. To inhibit autophagy, cells were treated with 250 μM chloroquine (Sigma) in complete growth medium for 4 hours. Untreated cells were incubated in medium containing an equal volume of DMSO. Following treatment, cells were rinsed with ice-cold PBS and whole cell protein extracts were collected in RIPA buffer. Flux was calculated as the increase in LC3-II levels in chloroquine-treated cells relative to untreated cells as determined through immunoblot.

### 2.15 Statistical analysis

Volcano plot and bubble charts were created using MATLAB_R2024a. All other graphs were made using GraphPad Prism v10.4.1. Imaging analysis was performed using ImageJ v1.54g. Hypothesis testing for normal distribution and equal variance was performed utilizing MATLAB_R2024a. Statistical significance was determined via GraphPad Prism v10.4.1 through unpaired *t-*test, unpaired *t-*test with Welch’s correction, or Mann-Whitney test, depending on hypothesis testing. Relevant statistical tests are indicated in figure legends.

## 3 RESULTS

### 3.1 Mammary fibroblasts secrete EVs in response to IR yet remain viable and are non-apoptotic

To study the effects of IR on the breast stromal microenvironment, we utilized human telomerase immortalized primary human breast fibroblasts isolated from reduction mammoplasty tissue, termed reduction mammary fibroblasts (RMFs) (Kuperwasser et al., 2004). Normal fibroblasts *in vitro* have been shown to produce substantially fewer vesicles than other cell types, such as cancer cells (Balaj et al., 2011). In our model, isolation methods that have been reported to produce highly pure EVs (Van Deun et al., 2014) resulted in insufficient sample material for repeated analyses. As a result, we implemented the isolation technique of differential ultracentrifugation (dUC), which remains the most-widely practiced method for EV isolation (Royo et al., 2020; S. Williams et al., 2023). RMFs were cultured for 48 hours immediately after exposure to IR followed by dUC isolation of EVs (**Figure 1a**). Western blot analysis confirmed the presence of common markers associated with EVs in our preparations. Particularly, tumor susceptibility gene 101 (TSG101) and syntenin-1 (Syn-1) (**Figure 1b**), and CD63 (**Figure S1a**), all of which are recommended markers according to the Minimal Information for the Study of Extracellular Vesicles (MISEV) guidelines (Théry et al., 2018; Welsh et al., 2024), were readily detected in our preparations. However, we were unable to detect GM130, a marker for Golgi-derived vesicles, in our EV preparations (**Figure S1b**), suggesting minimal or no contamination from intracellular organelles. Furthermore, transmission electron microscopy imaging of EV samples demonstrated the presence of vesicular structures (**Figure S1c**). Together, these data confirm that the UC-based procedure implemented in our model results in the sufficient isolation of EVs.

Nanoparticle tracking analysis (NTA) of EVs from irradiated and control RMFs revealed nearly identical size distributions, with a peak diameter of approximately 150 nm (**Figure 1c**), while particle concentrations were substantially elevated in the cases where RMFs were irradiated. When exposed to an IR dose of 10 Gy, RMFs secreted approximately 3.4x the concentration of EVs compared to the control (**Figure 1d**). RMFs treated to intermediate IR doses (2 and 6 Gy) secreted approximately 2x the number of EVs compared to the control. Importantly, EV samples from both control and irradiated RMFs were highly susceptible to detergent-mediated disruption, indicating a large concentration of membranous particles (Jeppesen et al., 2019) (**Figure S1d**). For the remainder of our studies, 10 Gy was utilized as dose for the irradiated condition.

Cellular exposure to IR damages genomic DNA both directly and indirectly, leading to DNA double-stranded breaks (Hall & Giaccia, 2012). Sustained DNA damage leads to cell cycle arrest while irreparable lesions typically induce cell death through apoptosis or mitotic catastrophe (Santivasi & Xia, 2014). Apoptosis as a mode of radiation-induced cell death is highly cell-type dependent. Previous studies have shown that fibroblast exposure to radiation results in cellular senescence but not apoptosis (Berzaghi et al., 2021; Papadopoulou & Kletsas, 2011). However, these studies have been conducted utilizing tumor-derived, cancer-associated fibroblasts (CAFs) or fibroblasts derived from organs other than the breast. Therefore, we sought to evaluate the cellular response with respect to cell survival and induction of apoptosis in our mammary fibroblast model. Viability was unchanged 48 hours after IR exposure with ≥ 95% live cells in control and irradiated conditions, across all experiments (**Figure 1e**). Additionally, cells were labelled with a fluorescent conjugate of Annexin V to assess the induction of apoptosis via flow cytometry. This analysis revealed no significant differences in the percentage of apoptotic cells when comparing irradiated and control cells (**Figure 1f**; **Figure S2**).

### 3.2 EVs derived from irradiated mammary fibroblasts contain damage associated molecular patterns

Next, we examined the cargo components associated with irradiated RMF EVs, with a focus on the proteomic profile. We implemented a tandem mass tag-based MS strategy to quantitatively characterize the proteins present within EVs from control and irradiated RMFs.

Our proteomic analysis consisted of duplicate samples for control and irradiated groups, each of which were TMT-labelled and combined. Through this approach, a total of 1,914 proteins were quantified (**Table S3**). We chose a 1.5-fold expression change (absolute value log_2_ ratio ≥ 0.59) and adjusted *p*-value < 0.05 as a cutoff for significance in irradiated RMF EVs. Based on these criteria, 36 proteins (34 upregulated, 2 downregulated) were significant (**Figure 2a**). Due to the limited number of significant downregulated proteins resulting from this analysis, we focused on the upregulated proteins. Interestingly, core canonical and linker histone variants predominated the list of upregulated proteins, with the top 5 proteins by fold-change all corresponding to histones, and an additional 4 histone variants in the remaining list of upregulated proteins (**Figure 2b**).

**Figure 2.**
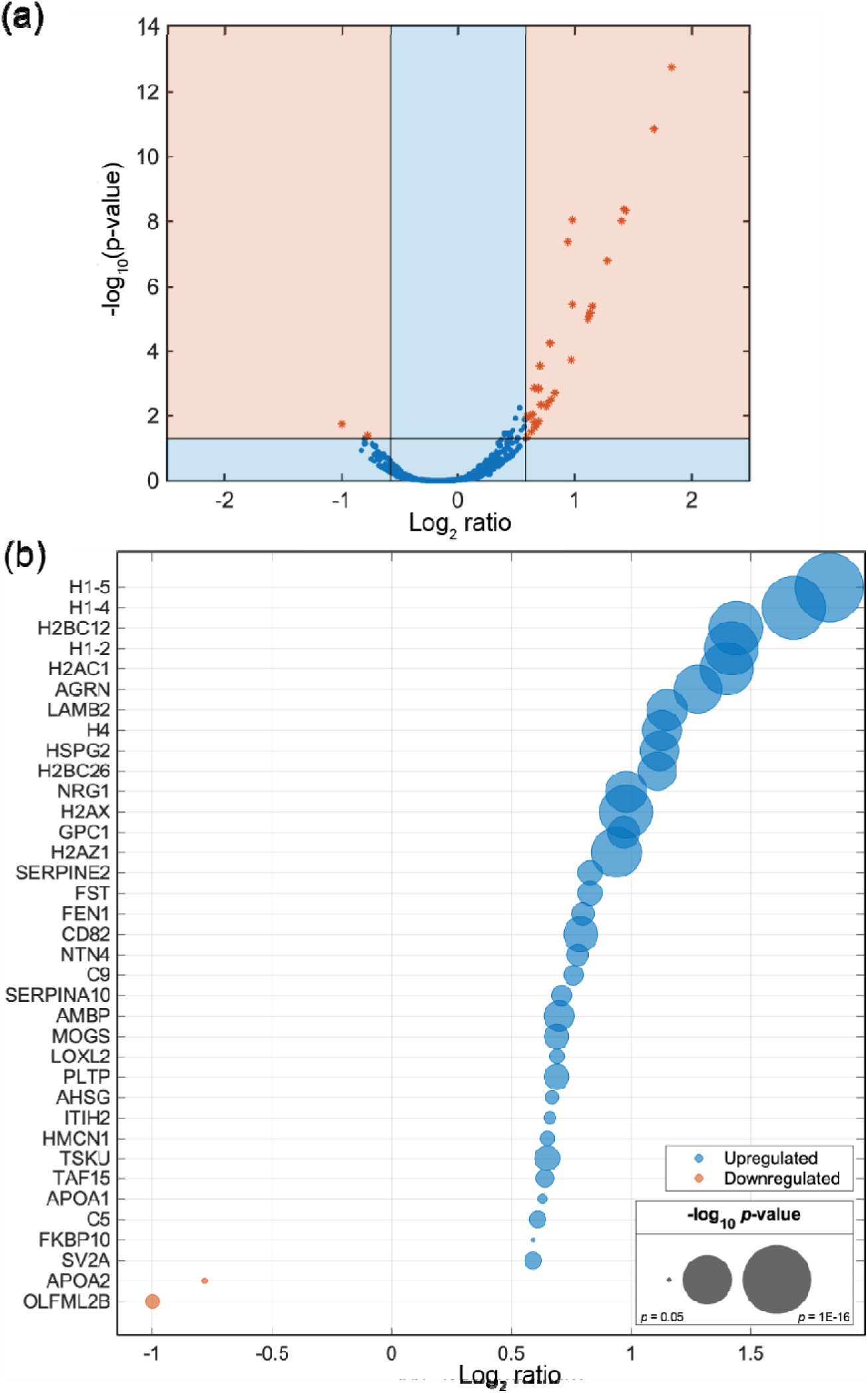
MS analysis of EVs derived from irradiated RMFs. **(a)** Volcano plot showing significant proteins (2 downregulated, 34 upregulated) in irradiated (IR; 10 Gy) *vs.* control (Ctrl; 0 Gy) RMF EVs. **(b)** Significant proteins determined through MS. X-axis quantification represents log_2_ ratio of IR / Ctrl. Bubble size represents −log_10_ adjusted *p*-value.

To systematically characterize and analyze the proteins upregulated in irradiated RMF EVs, we performed (GO) enrichment analysis on the significant upregulated proteins (**Table S4**). The proteins upregulated in irradiated RMF EVs largely represented factors indicative of a damage response. Injured tissues release damage associated molecular patterns (DAMPs) from intracellular and extracellular compartments that activate the innate immune system (Vénéreau et al., 2015). DAMPs, however, do not necessarily originate from dead cells and can be secreted or exposed by living cells undergoing a stress response, providing a mechanism for rapidly alerting the immune system. DAMPs autonomously trigger inflammatory processes through interaction with pattern recognition receptors (PRRs) on both immune and non-immune cells (Liu & Cao, 2016). Proteins that we identified as extracellular and intracellular DAMPs based on UniProt and/or GO annotations (**Table 1**), and those reported as having wound healing or inflammation-related functions (**Table 2**) were manually selected from our upregulated protein list. Our analysis revealed that irradiated RMF EVs are replete with intracellular DAMPs, consisting primarily of nucleosome proteins and heterochromatin components (**Figure S3a**), which when presented in cytosolic or extracellular contexts trigger a potent immune response (Richards et al., 2022). Additionally, we identified many extracellular matrix (ECM) proteins upregulated in irradiated RMF EVs, particularly basement membrane components (**Figure S3b**), which function as extracellular DAMPs when fragmented or solubilized and presented outside of their native context and structure (Gupta et al., 2022). The remaining list (**Table 2**) consists of molecules that are related to a DAMP response (hereinafter termed *DAMP-related molecules*), with negative regulation of hydrolase activity (GO:0051346), negative regulation of signal transduction (GO:0009968), and negative regulation of response to stimulus (GO:0048585) indicated as the top biological process terms (**Table S4**). Also of note, these DAMP-related molecules seem to be associated more specifically with EV-based secretion (**Figure S3c**).

**Table 1.**
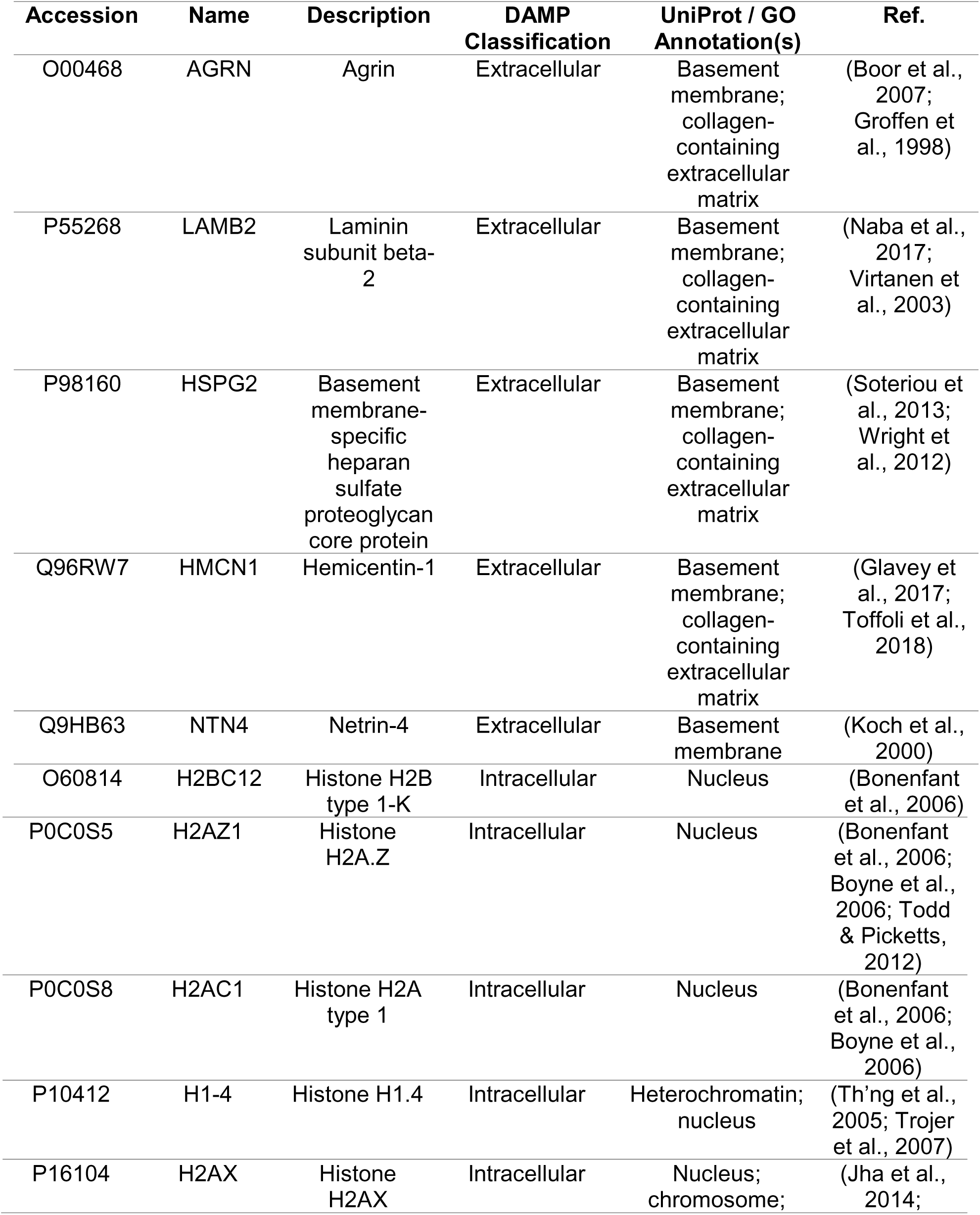
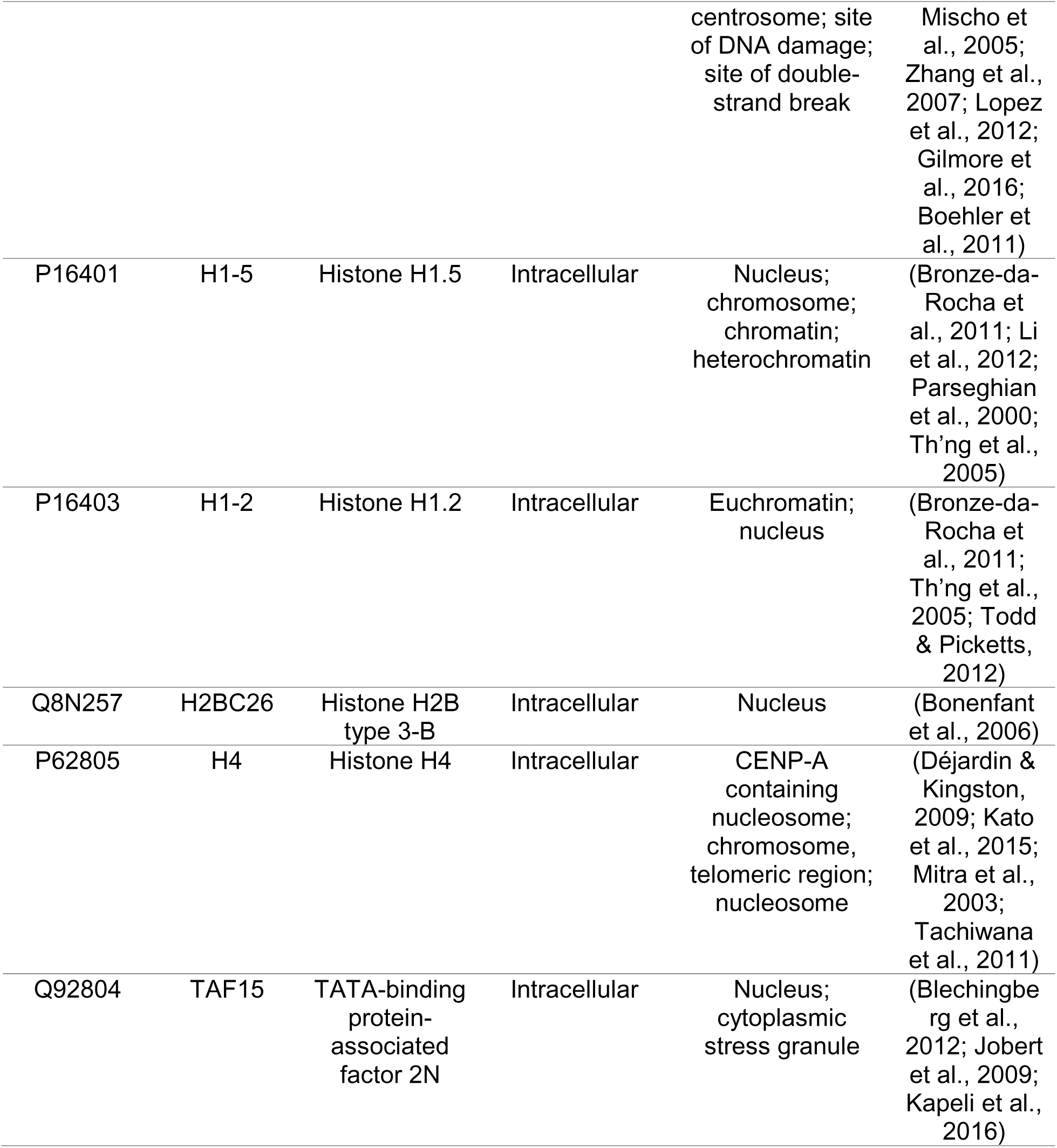
Extracellular and intracellular DAMPs identified in significant upregulated proteins in irradiated RMF EVs.

| Accession | Name | Description | DAMP Classification | UniProt / GO Annotation(s) | Ref. |
| --- | --- | --- | --- | --- | --- |
| O00468 | AGRN | Agrin | Extracellular | Basement membrane; collagen-containing extracellular matrix | (Boor et al., 2007; Groffen et al., 1998) |
| P55268 | LAMB2 | Laminin subunit beta-2 | Extracellular | Basement membrane; collagen-containing extracellular matrix | (Naba et al., 2017; Virtanen et al., 2003) |
| P98160 | HSPG2 | Basement membrane-specific heparan sulfate proteoglycan core protein | Extracellular | Basement membrane; collagen-containing extracellular matrix | (Soteriou et al., 2013; Wright et al., 2012) |
| Q96RW7 | HMCN1 | Hemicentin-1 | Extracellular | Basement membrane; collagen-containing extracellular matrix | (Glavey et al., 2017; Toffoli et al., 2018) |
| Q9HB63 | NTN4 | Netrin-4 | Extracellular | Basement membrane | (Koch et al., 2000) |
| O60814 | H2BC12 | Histone H2B type 1-K | Intracellular | Nucleus | (Bonenfant et al., 2006) |
| P0C0S5 | H2AZ1 | Histone H2A.Z | Intracellular | Nucleus | (Bonenfant et al., 2006; Boyne et al., 2006; Todd & Picketts, 2012) |
| P0C0S8 | H2AC1 | Histone H2A type 1 | Intracellular | Nucleus | (Bonenfant et al., 2006; Boyne et al., 2006) |
| P10412 | H1-4 | Histone H1.4 | Intracellular | Heterochromatin; nucleus | (Th'ng et al., 2005; Trojer et al., 2007) |
| P16104 | H2AX | Histone H2AX | Intracellular | Nucleus; chromosome; | (Jha et al., 2014; |
|  |  |  |  | centrosome; site of DNA damage; site of double-strand break | Mischo et al., 2005; Zhang et al., 2007; Lopez et al., 2012; Gilmore et al., 2016; Boehler et al., 2011) |
| P16401 | H1-5 | Histone H1.5 | Intracellular | Nucleus; chromosome; chromatin; heterochromatin | (Bronze-da-Rocha et al., 2011; Li et al., 2012; Parseghian et al., 2000; Th'ng et al., 2005) |
| P16403 | H1-2 | Histone H1.2 | Intracellular | Euchromatin; nucleus | (Bronze-da-Rocha et al., 2011; Th'ng et al., 2005; Todd & Picketts, 2012) |
| Q8N257 | H2BC26 | Histone H2B type 3-B | Intracellular | Nucleus | (Bonenfant et al., 2006) |
| P62805 | H4 | Histone H4 | Intracellular | CENP-A containing nucleosome; chromosome, telomeric region; nucleosome | (Déjardin & Kingston, 2009; Kato et al., 2015; Mitra et al., 2003; Tachiwana et al., 2011) |
| Q92804 | TAF15 | TATA-binding protein-associated factor 2N | Intracellular | Nucleus; cytoplasmic stress granule | (Blechingberg et al., 2012; Jobert et al., 2009; Kapeli et al., 2016) |

**Table 2.** DAMP-related molecules identified in significant upregulated proteins in irradiated RMF EVs.

|  | Name | Description | Relevant GO Annotation(s) | Ref. |
| --- | --- | --- | --- | --- |
| P01031 | C5 | Complement C5 | Chemokine activity; inflammatory response; negative regulation of macrophage chemotaxis; positive regulation of chemokine production | (Ambati et al., 2003; Floreani et al., 1998; Luu et al., 2000; Nilsson et al., 1996) |
| P02647 | APOA1 | Apolipoprotein A-I | Negative regulation of cytokine production involved in immune response; negative regulation of inflammatory response; negative regulation of interleukin-1 beta production; negative regulation of response to cytokine stimulus; negative regulation of tumor necrosis factor-mediated signaling pathway; positive regulation of phagocytosis | (Di Bartolo et al., 2011; Furlaneto et al., 2002; Tanaka et al., 2010) |
| P02748 | C9 | Complement component C9 | Positive regulation of immune response | (Menny et al., 2018) |
| P02760 | AMBP | Protein AMBP | Antioxidant and tissue repair; Negative regulation of inflammatory response; endopeptidase inhibitor activity; negative regulation of JNK cascade | (Berggård et al., 1999; Jersmann et al., 2003; Kobayashi et al., 2003) |
| P02765 | AHSG | Alpha-2-HS-glycoprotein | Regulation of inflammatory response; positive regulation of phagocytosis; endopeptidase inhibitor activity; pinocytosis | (Jersmann et al., 2003; Ketteler et al., 2003; Lord, 2003) |
| P07093 | SERPINE2 | Glia-derived nexin | Serine-type endopeptidase inhibitor activity; heparin binding; regulation of cell migration | (Bouton et al., 2007; Evans et al., 1991; Gloor et al., 1986; Koistinen et al., 2009) |
| P19823 | ITIH2 | Inter-alpha-trypsin inhibitor heavy chain H2 | endopeptidase inhibitor activity; hyaluronic acid binding | (Enghild et al., 1989; Sanggaard et al., 2010) |
| P19883 | FST | Follistatin | TGβ-family antagonist; heparan sulfate proteoglycan binding | (Chen et al., 2003; |
|  |  |  |  | Schneyer et al., 2003) |
| P27701 | CD82 | CD82 antigen | Regulation of inflammatory response | (Lee et al., 2023) |
| P35052 | GPC1 | Glypican-1 | Heparan sulfate proteoglycan catabolic process; negative regulation of fibroblast growth factor receptor signaling pathway | (Gutiérrez & Brandan, 2010; Mani et al., 2003) |
| P39748 | FEN1 | Flap endonuclease 1 | DNA binding; damaged DNA binding; DNA-repair; endonuclease activity; exonuclease activity | (Harrington & Lieber, 1994; Hosfield et al., 1998; Murray et al., 1994; Qiu et al., 2002) |
| Q02297 | NRG1 | Pro-neuregulin-1, membrane-bound isoform | Cytokine activity; cell communication; wound healing | (Britsch, 2007; Esper et al., 2006; Osheroff et al., 1999) |
| Q8WUA8 | TSKU | Tsukushin | Negative regulation of myofibroblast differentiation; negative regulation of TGFβ-1 production; negative regulation of Wnt signaling pathway; wound healing | (Niimori et al., 2014; Ohta et al., 2011) |
| Q96AY3 | FKBP10 | Peptidyl-prolyl cis-trans isomerase FKBP10 | Collagen fibril organization; extracellular matrix assembly; wound healing | (Liang et al., 2017; Lietman et al., 2014, 2017) |
| Q9Y4K0 | LOXL2 | Lysyl oxidase homolog 2 | Collagen fibril organization; positive regulation of epithelial to mesenchymal transition; endothelial cell migration; endothelial cell proliferation | (Bignon et al., 2011; Millanes-Romero et al., 2013) |
| P55058 | PLTP | Phospholipid transfer protein | Negative regulation of inflammatory response; negative regulation of macrophage cytokine production | (Vuletic et al., 2011) |

While extracellular and intracellular DAMPs, such as those listed in **Table 1**, would canonically induce inflammatory processes through PRR activation, some additional functional enrichments were revealed through GO analysis. Particularly for the extracellular DAMPs, basement membrane organization (GO:0071711) and extracellular matrix assembly (GO:0085029) were enriched biological process terms (**Table S4**). Therefore, some proteins secreted through EVs may elicit an inflammatory response through PRR recognition while others may regulate other key biological processes as part of a wound healing response. Fibroblasts are well known to undergo activation involving cell recruitment and migration as well as remodeling of the ECM in response to tissue damage (Cialdai et al., 2022). Thus, EV-associated proteins secreted in response to IR may play a crucial role in these processes. However, basement membrane assembly is not known to be facilitated by EVs (Hellicar et al., 2022; McCaughey & Stephens, 2019), although basement membrane proteins have been identified in EVs isolated from patient-derived biofluids and *in vitro* models (Gonzalez-Begne et al., 2009; Kistenmacher et al., 2024; McKay et al., 2020; Principe et al., 2013).

### 3.3 Irradiated mammary fibroblasts display hallmark characteristics of senescence yet secrete a unique subset of proteins through EVs

To further investigate the phenotypic development associated with enhanced EV secretion in irradiated RMFs, we examined the induction of a senescence-like phenotype following acute exposure to IR. It is known that a single dose of γ-radiation (*i.e*., 10 Gy) is sufficient to induce senescence after 7–10 days *in vitro* (Kohli et al., 2021). However, the course of development of the senescent phenotype is rarely studied. Furthermore, it is well known that exposure to ionizing radiation activates the DNA-damage response (DDR), subsequently activating p53 and leading to growth arrest through induced expression of the cell cycle regulator p21^cip1/waf1^ (He et al., 2005). In our model, DDR activation was observed in irradiated RMFs, measured through the detection of phosphorylated H_2_AX foci (γH_2_AX; **Figure 3a**). Additionally, p53 expression was transiently elevated in RMFs following exposure to IR, with a return to baseline after 24 hours (**Figure S4a**). In contrast, the expression of p21 was sustained at 48 hours, indicating that irradiated RMFs became growth-arrested (**Figure S4b**). To further examine the degree of growth arrest, we incubated RMFs with 5-Ethynyl-2’-deoxyuridine (EdU). Incorporation was observed in control RMFs (approx. 60% of cells) yet was rare in irradiated cells (approx. 5% of cells), revealing that RMFs are no longer dividing 48 hours post-IR (**Figure 3b**). Additionally, senescence associated β-galactosidase (SA-βgal) activity was detected in irradiated cells (**Figure 3c**).

**Figure 3.**
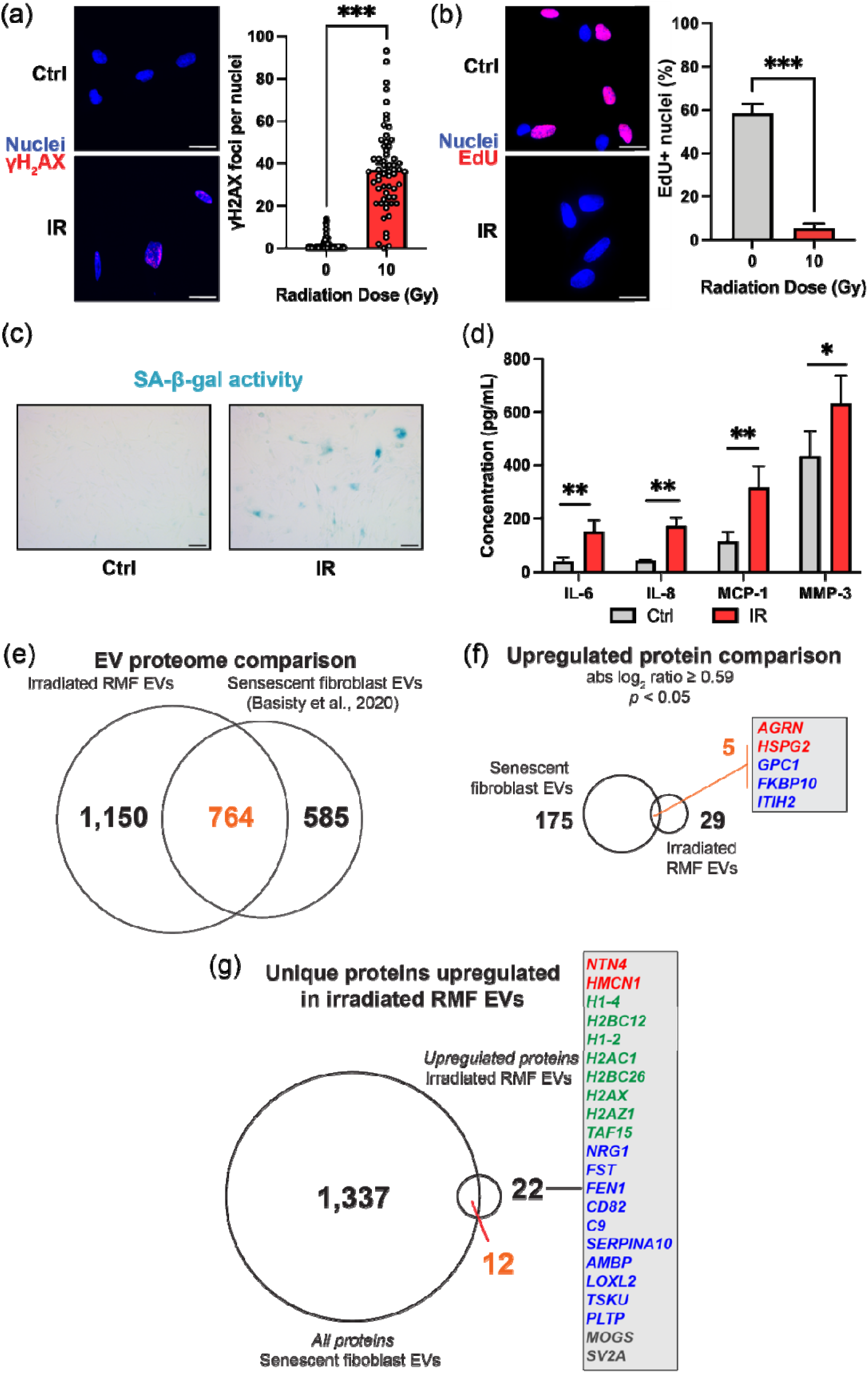
Irradiated RMFs display hallmark characteristics of senescence yet secrete a unique subset of proteins through EVs. **(a)** Representative fluorescence microscopy images of control (Ctrl; 0 Gy) and irradiated (IR; 10 Gy) RMFs fixed 30 min post-IR and stained for phosphorylated histone H2A.X (γH_2_AX, red). Nuclei are visualized in blue. Quantification represents γH_2_AX foci per cell (N=3). Scale bar represents 25 μm. Statistical analysis was performed using Mann-Whitney test. **(b)** Representative fluorescence microscopy images of 5-ethynyl-2’-deoxyuridine (EdU) incorporation in control and irradiated RMFs. Quantification represents percentage of nuclei (blue) positive for EdU (red) (N=3). Scale bar represents 25 μm. Statistical analysis was performed using Mann-Whitney test. **(c)** Representative brightfield microscopy images of senescence-associated β-galactosidase (SA-β-gal) activity (blue) in control and irradiated RMFs. Scale bar represents 100 μm**. (d)** Concentration (pg/mL) of indicated proteins in the CM of control and irradiated RMFs, determined through Luminex assay (N=3). Statistical analysis was performed using unpaired *t*-tests **(e)** Venn diagram comparing all proteins quantified in EVs secreted by RMFs 48 hours post-IR and senescent human fibroblasts (data from SASP atlas; www.saspatlas.org; Basisty *et al.,* 2020). **(f)** Venn diagram comparing significant upregulated proteins in irradiated RMF EVs and senescent fibroblast EVs. Proteins listed in inset were upregulated in both conditions and are colored according to their corresponding damage associated molecular pattern (DAMP) classification (red=extracellular DAMP, blue=DAMP-related molecule). **(g)** Venn diagram comparing upregulated proteins in irradiated RMF EVs and all proteins identified in senescent fibroblast EVs. Proteins listed in inset are unique to the upregulated proteome of irradiated RMF EVs and are colored according to their classification (red=extracellular DAMP, green=intracellular DAMP, blue=DAMP-related molecule, gray=other). Statistical significance was determined as described with \**p* < 0.05, \*\**p* < 0.01, and \*\*\**p* < 0.001. Error bars show standard deviation.

In addition to growth arrest, senescent cells are understood to develop a heterogenous secretory program generally termed the senescence-associated secretory phenotype (SASP) (Coppé et al., 2008). Thus, we also probed for the presence of relevant SASP factors in the conditioned media (CM) of irradiated RMFs 48 hours post-IR. Due to the heterogeneity of the SASP, our analysis focused on a fibroblast-relevant response, probing for key pro-inflammatory cytokines as well matrix remodeling components, including matrix metalloproteinases (MMPs) and tissue inhibitors of metalloproteinases (TIMPs) (**Table S1–2**; **Figure S4c**). Several cytokines were detected at significantly elevated levels in the CM of irradiated RMFs (**Figure 3d**). Notably, IL-6 (3.8x), IL-8 (3.9x), and MCP-1 (2.8x) were all elevated post-IR. Additionally, MMP-3 was elevated at 1.5x in irradiated RMF CM.

In addition to secretion of soluble proteins, there is evidence that EV secretion is a component of the SASP (Wallis et al., 2020). Recently, the secretome of senescent primary human fibroblasts (IMR-90) was evaluated (Basisty et al., 2020). In their study, multiple senescence-inducing agents were utilized, and the proteomes of isolated EVs were analyzed, creating the SASP Atlas database (http://saspatlas.com). We leveraged this database by comparing the results of the proteomic analysis of IR-induced EV secretion from our study (irradiated RMF EVs) to the fibroblast proteome from the SASP Atlas (senescent fibroblast EVs). For a direct comparison, we examined only IR-induced senescence data. Importantly, the same dUC-based EV isolation procedure was conducted in both studies. In our analysis, we found that more proteins were quantified in irradiated RMF EVs than in senescent fibroblasts EVs (1,914 *vs.* 1,349) (**Figure 3e**). The majority of proteins present in senescent fibroblast EVs (764 / 1,349; approx. 57%) were also identified in irradiated RMF EVs. However, among the 180 proteins that were significantly upregulated in senescent fibroblast EVs (log_2_ ratio ≥ 0.59, *p* < 0.05) only 5 were in common with upregulated proteins from irradiated RMF EVs (**Figure 3f**): agrin (AGRN), perlecan (HSPG2), glypican-1 (GPC1), inter-alpha-trypsin inhibitor heavy chain H2 (ITIH2), and FK506 binding protein 10 (FKBP10). Notably, 4 proteins that were upregulated in irradiated RMF EVs were in fact downregulated in senescent fibroblast EVs: histone H1.5 (H1-5), histone H4 (H4), apolipoprotein A1 (APOA1), and complement C5 (C5) (**Figure S4d**). Also, 22 of the upregulated proteins in irradiated RMF EVs were not identified at all in senescent fibroblast EVs (**Figure 3g**). This collection of proteins may represent distinct cargo components that are uniquely secreted as part of an acute response to IR in mammary-derived fibroblasts.

### 3.4 DAMP release through EVs in irradiated mammary fibroblasts is dependent on Rab27a

The canonical cellular response to IR-induced damage involves p53 activation, which may lead to downstream growth arrest and senescence (Suzuki et al., 2001). To study the contribution of these key signaling pathways on the acute EV secretion response to IR in mammary fibroblasts, we first probed the effect of p53 activation in the absence of IR exposure. Basal levels of p53 are maintained through negative regulation by the E3 ubiquitin ligase MDM2 (Momand et al., 1992). Thus, we treated fibroblasts with Nutlin-3a, a small molecule inhibitor of MDM2. As expected, incubation of RMFs in Nutlin-3a-containing growth medium resulted in a substantial elevation in p53 levels (**Figure S5a**). Following 72 hour treatment with Nutlin-3a, EVs were isolated, and size and concentration of EVs were determined through NTA. EVs from Nutlin-3a-treated cells had a similar size distribution to that of control and irradiated EVs, with a peak diameter of 150 nm (**Figure S5b**). Particle concentrations were substantially elevated in the case of RMFs treated with Nutlin-3a (**Figure 4a**)—an approximate 7x increase in secretion rate compared to that of the control., indicating a very pronounced EV secretion in response to p53 activation. Of note, the level of p53 expression in RMFs following Nutlin-3a treatment was similar to that of irradiated RMFs (**Figure S4c**). However, in our IR model, p53 activation was shown to be transient, with a return to baseline occurring within 24 hours. This activation impulse is seemingly insufficient in eliciting the same regulatory impact on downstream effector genes as Nutlin-3a. Thus, p53 activation alone may not be the primary driver of enhanced EV secretion in irradiated RMFs. To further probe the role of p53 activation on the altered EV secretion phenotype in irradiated RMFs, we conducted an shRNA knockdown (KD) of p53 (**Figure S5c**) then exposed cell cultures to IR and isolated EVs from CM. Unexpectedly, irradiated p53 KD RMFs secreted 2.7x more EVs than unirradiated p53 KD cells (**Figure 4b**). These combined results indicate that post-IR p53 activation contributes to enhanced EV secretion but is largely not responsible for the several-fold increase in EV concentration.

**Figure 4.**
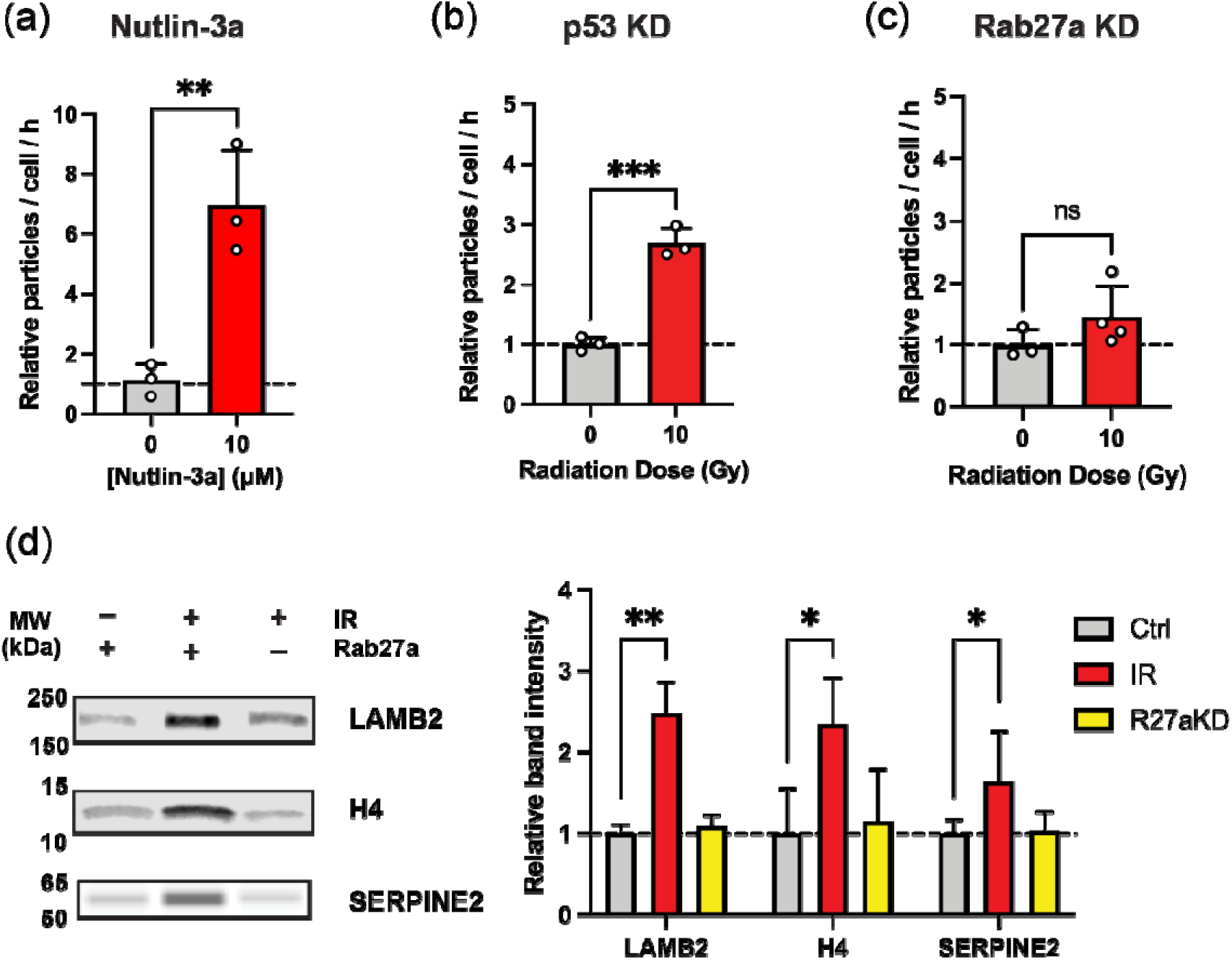
Secretion of key DAMPs through EVs by irradiated RMFs is dependent on Rab27a. **(a)** Particle concentration normalized to cell number per hour for EVs derived from RMFs treated with Nutlin-3a for 72 hours (N=3), determined via NTA. **(b)** Particle concentration normalized to cell number per hour for EV preparations from control (Ctrl; 0 Gy) and irradiated (IR; 10 Gy) p53 KD and **(c)** Rab27a KD RMFs, determined via NTA (N=3). **(d)** Representative western blot analysis of selected DAMPs—LAMB2, Histone H4, and SERPINE2—in EV-derived protein extracts from indicated experimental conditions. Quantification shows relative band intensity normalized to total protein stain (N=3). Statistical significance was determined via unpaired *t-*tests, with nonsignificant (ns) *p* > 0.05, \**p* < 0.05, \*\**p* < 0.01, and \*\*\**p* < 0.001. Error bars show standard deviation.

Since p53 activation was not the key mechanism in our model, we sought to determine the contribution of intracellular vesicle secretion machinery on enhanced EV secretion in irradiated RMFs. Ras-associated binding proteins (Rab) are small GTPases localized in distinct membrane-bound compartments. The importance of Rab proteins has been highlighted in the regulation of vesicle trafficking processes including vesicle formation, motility, tethering and fusion to the acceptor membrane, and signaling to other organelles (Corbeel & Freson, 2008). Several Rab proteins play major roles EV secretion (Blanc & Vidal, 2018). In our proteomics analysis, 20 Rab family proteins, including Rab27a, were identified in RMF EVs (**Table S5**). Rab27, which regulates the transport and docking of multivesicular bodies (MVBs) to the plasma membrane (Van Niel et al., 2018), is by far the most well-studied Rab protein in EV research. We therefore focused on the contribution of Rab27a in the EV-associated response to IR. We conducted a KD of Rab27a in RMFs (**Figure S5d**), which did not affect EV size distribution (**Figure S5b**). Surprisingly, Rab27a KD abrogated the IR-induced enhanced secretion of EVs in RMFs (**Figure 4c**). These results suggest a direct dependence on Rab27a mechanistically in the enhanced secretion of EVs in irradiated RMFs.

To further probe the influence of Rab27a in the secretion of specific cargoes through EVs, we performed western blot analysis on EVs derived from irradiated Rab27a KD RMFs. We focused on proteins from our MS analysis that were representative of the 2 classes of DAMPs as well as the subset of DAMP-related molecules. As such, we selected the proteins laminin subunit β-2 (LAMB2), histone H4, and serpin family E member 2 (SERPINE2) to represent extracellular DAMPs, intracellular DAMPs, and DAMP-related molecules, respectively. We confirmed the significant upregulation and quantitative results from our MS analysis for all three proteins through comparison of EVs derived from unirradiated and irradiated scramble control RMFs (**Figure 4d**; **Table 3; Figure S6**). Interestingly, EVs derived from irradiated Rab27a KD RMFs expressed comparable levels of all three proteins to EVs derived from unirradiated scramble control RMFs. These results demonstrate that both the enhanced EV secretion and enrichment of DAMPs observed in irradiated RMFs are dependent on Rab27a. Of note, a recent study reported a dependence on Rab27a for lysosomal secretion in senescent cells, involving the induction of macroautophagy (Rovira et al., 2022). However, we did not observe an increase in macroautophagy post-IR (**Figure S7**). While a more comprehensive analysis of lysosomal dynamics may be needed, these combined data suggest that Rab27a-mediated MVB trafficking drives the acute response to IR, resulting in DAMP-containing EV secretion.

**Table 3.**
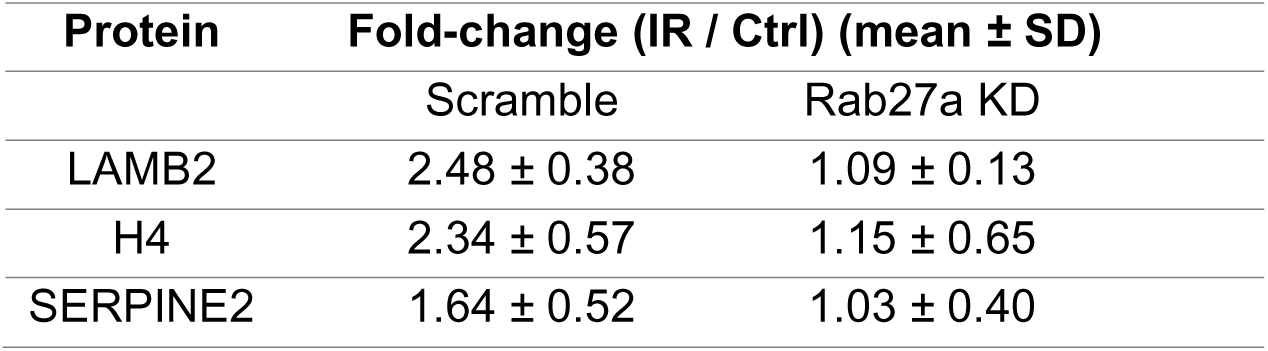
Summary of quantitative results from Western blot analysis of selected DAMPs in EV-derived protein extracts from irradiated scramble control and Rab27a KD RMFs.

## 4 DISCUSSION

Despite aggressive therapy, breast cancer patients, particularly those with triple-negative breast cancer, are at high risk for recurrence (Adkins et al., 2011). The impact of the stromal response to IR and the resulting microenvironmental alterations are critical for determining risk of recurrence following radiotherapy. Acute or early reactions to IR are characterized by rapidly occurring changes within hours to days of exposure (De Ruysscher et al., 2019). Early molecular changes, including activation of signaling pathways and induction of intercellular communication, occur shortly after IR exposure, whereas the cellular events and tissue remodeling processes triggered by these mechanisms evolve more slowly (Rodemann & Blaese, 2007). This suggests that processes such as radiation-induced cell death and apoptosis do not drive underlying late tissue reactions. Rather, the molecular and cellular processes responsible for tissue remodeling are triggered through alterations in cellular signaling.

Others have previously shown that cellular exposure to IR results in substantially enhanced EV production in models of head and neck (Mutschelknaus et al., 2016) and lung cancer (Yu et al., 2006). However, several other studies have shown the contrary—that exposure to IR does not impact EV secretion (Berzaghi et al., 2021; Colangelo & Azzam, 2020). Thus, current evidence suggests that the EV secretion response to IR is a cell type and context-dependent process. In our model, we report a substantial increase in release of EVs after acute exposure to IR. Exposure to IR can lead to irreparable genetic lesions, which may eventually result in apoptosis. Cells undergoing apoptosis release large amounts of EVs due to cell membrane budding and blebbing (Li et al., 2020). In our study, however, there was no measurable induction of cell death or apoptosis in irradiated mammary fibroblasts. Some studies have shown that p53 plays an important role in IR-induced EV secretion (Lespagnol et al., 2008; Yu et al., 2006). While our study corroborated findings of an enhancement in EV secretion as a result of p53 activation, this was not the primary driving force for post-IR EV secretion in our model. Rather, Rab27a-dependent MVB trafficking was shown to be the key mechanism.

Quantitative analysis of EVs from irradiated mammary fibroblasts revealed an unconventional proteomic profile. Notably, irradiated fibroblast EVs were highly enriched in histones. While histones are frequently identified in proteomic analyses of isolated EVs (H. Zhang et al., 2018), studies investigating the presence of histones within EVs have produced contradictory results. Several studies have claimed that histones are associated with or packaged within EVs (Lázaro-Ibáñez et al., 2019; Takahashi et al., 2017) while others claim that secretion of histones is not a part of classical exosome secretion and instead may result from technical artifacts or due to fusion of organelles (*i.e.* amphisome formation) (Jeppesen et al., 2019). Other recent work has shown that neutrophils explicitly package chromatin into MVBs, whereby histones and DNA can be secreted extracellularly through EVs (Arya et al., 2025). Additionally, in a landmark study using rat oligodendrocytes as a model, core and linker histones were identified to be secreted specifically via the MVB pathway (Singh et al., 2025). A robust and consistent upregulation of EV histone secretion was observed in response to various stress conditions, namely, heat stress, endotoxin exposure, hypoxia, and H_2_O_2_-induced oxidative stress. While these stress paradigms carry cellular and molecular nuance, including organism-level differences (Kumari, 2025; Wasserman et al., 2022), the current evidence points to a shared response involving histone secretion through EVs in mammalian cells. Indeed, the substantial upregulation of histones identified in our study of EV preparations derived from human cells treated with γ-radiation agrees with the current evidence of histone secretion in stress contexts.

Nonetheless, further characterization of the histone profile identified in our study is warranted, including analysis of membrane binding or association, luminal presence, and analysis of post translational modifications. The most compelling evidence to date suggests that EV-associated histones are mostly localized to the outer surface of EVs and lack extensive post-translational modifications (Singh et al., 2025). In the context of radiation-induced injury, this remains to be studied, as the molecular nature of EV-associated histones would dictate the intended biological consequences (Buzas, 2022). Histones are well-characterized as a major class of DAMPs following tissue injury or cell death (Roh & Sohn, 2018). Extracellular histones have been considered as potential mediators of lethal systemic inflammatory diseases including those involving infection and sterile inflammation (Chen et al., 2014). The secretion of histones through EVs may therefore allow radiation-injured cells to trigger a potent inflammatory response.

Beyond intracellular compartment proteins, extracellular proteins are typically components of the ECM such as glycoproteins and proteoglycans largely make up the second major class of DAMPs (Frevert et al., 2018). The ECM provides critical architectural structure in all tissues and creates a microenvironment for chemical and mechanical interactions that are crucial for tissue homeostasis (Nanini et al., 2018). However, during tissue injury, cleavage of ECM molecules produces protein fragments and exposes cryptic epitopes, which may activate and recruit immune cells (Lambert et al., 2021). We identified several ECM components in irradiated mammary fibroblast EVs, including agrin, a large heparan sulfate (HS) proteoglycan, and LAMB2, both of which are major constituents of the basement membrane (Naylor et al., 2020). Under physiologic conditions, proteoglycans are not soluble. However, stimulation with soluble agrin has been reported to activate the MAPK pathway in macrophages (Mazzon et al., 2012) and positively modulate activation in T cells (Kabouridis et al., 2012; Khan et al., 2001). Additionally, HS chains themselves directly activate several PRRs (McQuitty et al., 2020). Interestingly, GPC1, a secreted glycoprotein with proteoglycan catabolic activity (Mani et al., 2003), was also upregulated in irradiated RMF EVs, highlighting a route for the breakdown of proteoglycans and release of soluble HS into the extracellular space (Goodall et al., 2014). In addition to proteoglycans, laminins play key structural roles in the ECM and have been documented in impacting the function of immune cells (Florea et al., 2016; Savino et al., 2015). Almost all leukocyte subpopulations express specific laminin receptors, and various laminins have been shown to interact with macrophages and monocytes, leading to increased inflammatory cytokine and MMP production (Simon & Bromberg, 2017). Thus, the release of fragments, subunits, and soluble forms of ECM proteins through EVs represents a critical immunomodulatory aspect of the damage response post-IR.

Beyond the two major classes of DAMPs, EVs derived from our irradiated fibroblast model contained a collection of proteins related to a DAMP response (Yamaguchi et al., 2020). Particularly, the remaining subset of proteins regulate macromolecular processes, with functions including enzyme inhibition, cytokine and chemokine activity, and the modulation of cellular signaling and inflammation. For example, complement system proteins C5 and C9 were both upregulated, which combine to form the membrane attack complex (MAC). While conventionally involved in the lysis of invading pathogens (Xie et al., 2020), recent studies have highlighted regulation of the innate immune response by the MAC in sterile inflammation. In fact, induction of signaling cascades and pro-inflammatory cytokine production has been described in human macrophages in response to MAC stimulation (Diaz-del-Olmo et al., 2021; Jimenez-Duran et al., 2022). Additionally, the serine protease inhibitor SERPINE2 was found to be upregulated in irradiated mammary fibroblast EVs in our study. Extracellular proteases and their inhibitors are critical in remodeling of the ECM (Mason & Joyce, 2010). Furthermore, SERPINE2 is upregulated in breast cancer (Fayard et al., 2009) and has been shown to be critical in metastasis, particularly through regulation of ECM density and macrophage polarization, although its role as an EV-associated protein has not been studied (Smirnova et al., 2016). Increased expression of SERPINE2 in radioresistant tumor cells has also been implicated in the lung cancer radiation response (J. Zhang et al., 2022). Thus, secretion of DAMP-related proteins through EVs could enable injured cells to communicate with their microenvironment through proteins that regulate cell signaling, inflammation, and ECM organization.

Fibroblasts have been described as having a robust and potent SASP response (Schafer et al., 2020). In our study, mammary fibroblasts adopted key characteristics of the SASP, reinforcing that even with acute exposure to IR, the breast stromal microenvironment may become highly altered toward a pro-inflammatory niche. Critically, MCP-1 chemokine secretion was elevated in our model, suggesting a response intended to recruit monocytes and macrophages. We have previously shown that macrophage infiltration of irradiated mammary tissue is an early step post-IR and is critical for CTC invasion *in vivo* (Rafat et al., 2018). Here, we show that IR-induced cytokine and chemokine secretion in mammary fibroblasts occurs in coordination with EV-associated secretion of DAMPs and related molecules, which could represent an integrated mechanism to facilitate the establishment of a pro-inflammatory and pro-wound healing microenvironment. While we showed that mammary fibroblasts secrete SASP factors post-IR, our study determined that the secretome of irradiated fibroblasts is distinct from that of cells in a later stage of senescence. Our proteomic comparison to senescent fibroblast-derived EVs revealed a unique subset of proteins that were enhanced in irradiated fibroblasts. While the precise function of this collection of proteins remains to be elucidated, it may represent a proteomic profile containing signals and inflammatory triggers that are crucial for directing cellular phenotypes in the breast post-IR.

This work begins to highlight key cellular and molecular alterations in the irradiated breast microenvironment through use of a model of normal breast stroma. Our study isolates the effects of IR on acute alterations in EV secretion in fibroblasts, a major cell type of the human breast stroma relevant to wound healing. How irradiated fibroblasts communicate with other cell types, in particular macrophages and other immune cells, is yet to be uncovered. Nonetheless, we report a novel alteration of the EV secretion profile in the early cellular response to IR that may have critical implications for the development of a pro-inflammatory breast tissue microenvironment post-radiotherapy.

## CONCLUSIONS

In this study, we examined the alterations in EV secretion in a model of breast stromal exposure to IR. We show that mammary fibroblasts adopt a senescence-like phenotype in the early stages of the radiation response, along with a novel reporting of various classes of DAMPs and related molecules associated with secreted EVs. Investigating EV cargo and function are critical in advancing our understanding of intercellular communication post-IR and may lead to improved techniques and methodologies to refine the application of IR in clinical settings.

## DATA AVAILABILITY STATEMENT

The data that support the findings of this study have been deposited to the ProteomeXchange Consortium via the PRIDE partner repository with the dataset identifier PXD066811.

## ACKNOWLEDGEMENTS

We thank Dr. Renee Dawson and the Vanderbilt Center for EV Research for equipment usage, Dr. Michael Freeman (Vanderbilt University Medical Center) for irradiator usage, Dr. Charlotte Kuperwasser (Tufts University) for providing immortalized reduction mammary fibroblasts, Dr. Evan Krystofiak (Vanderbilt Cell Imaging Shared Resource) for assistance on electron microscopy sample preparation, Dr. James McBride and the Vanderbilt Institute for Nanoscale Science and Engineering (VINSE) for electron microscopy imaging, the Proteomics Laboratory at the Vanderbilt Mass Spectrometry Research Center for support in MS experimentation, and former undergraduate student Akhila Ramgiri (Vanderbilt University) for early contributions. This research was financially supported by a VINSE pilot award, the Concern Foundation Conquer Cancer Now Award, the Vanderbilt Institute for Clinical and Translational Research grant no. V0000039983, the National Institutes of Health under awards R00CA201304 and T32DK101003, and the National Science Foundation Graduate Research Fellowship Program under grant nos. 1937963 and 2444112. This work includes schematics created using BioRender.

## AUTHOR CONTRIBUTIONS

**Greg Berumen Sánchez:** conceptualization, methodology, software, validation, formal analysis, investigation, data curation, writing–original draft, writing–review and editing, visualization, funding acquisition. **Purvi Patel:** investigation, validation, writing–review and editing. **Preston Gomez-Crase:** investigation. **Chloe Kim:** investigation. **Kristie Lindsey Rose:** methodology, validation, formal analysis, resources, data curation, writing–review and editing, project administration. **Marjan Rafat:** conceptualization, methodology, resources, writing–review and editing, supervision, project administration, funding acquisition.

## ABBREVIATIONS

CAF: cancer-associated fibroblast
CM: conditioned media
CTC: circulating tumor cell
CV: Coefficient of variation
DAMP: damage associated molecular pattern
DDR: DNA damage response
DMEM: Dulbecco’s modified eagle medium
DMSO: dimethyl sulfoxide
DPBS: Dulbecco’s phosphate buffered saline
ECM: extracellular matrix
EdU: 5-ethynyl-2’-deoxyuridine
EV: extracellular vesicle
GO: gene ontology
HCD: higher-energy collisional dissociation
HEPES: 4-(2-hydroxyethyl)-1-piperazineethanesulfonic acid
HRP: horseradish peroxidase
HS: heparan sulfate
IL: interleukin
IFN: interferon
IR: ionizing radiation
KD: knock down
LC3: microtubule-associated protein 1A/1B-light chain 3
MAC: membrane attack complex
MAPK: Mitogen-Activated Protein Kinase
MCP: monocyte chemoattractant protein
MISEV: minimal information for the study of extracellular vesicles
MMP: matrix metalloproteinase
MVB: multi-vesicular body
NTA: nanoparticle tracking analysis
PBS: phosphate-buffered saline
PES: polyether sulfone
PRR: pattern recognition receptor
PVDF: polyvinylidene fluoride
RIPA: radioimmunoprecipitation assay
RMF: reduction mammary fibroblast
RPM: revolutions per minute
SASP: senescence associated secretory phenotype
SA-βgal: senescence associated β-galactosidase
shRNA: short-hairpin RNA
STS: staurosporine
TBS: tris-buffered saline
TEM: Transmission electron microscopy
TIMP: tissue inhibitor of metalloproteinase
TMT: tandem mass tag
TNF: tumor necrosis factor
UC: ultracentrifugation

