## Supplementary Figures for "Mammary Fibroblasts Secrete Damage Associated Molecular Patterns through Extracellular Vesicles in Response to Ionizing Radiation"

**SUPPLEMENTAL FIGURES**

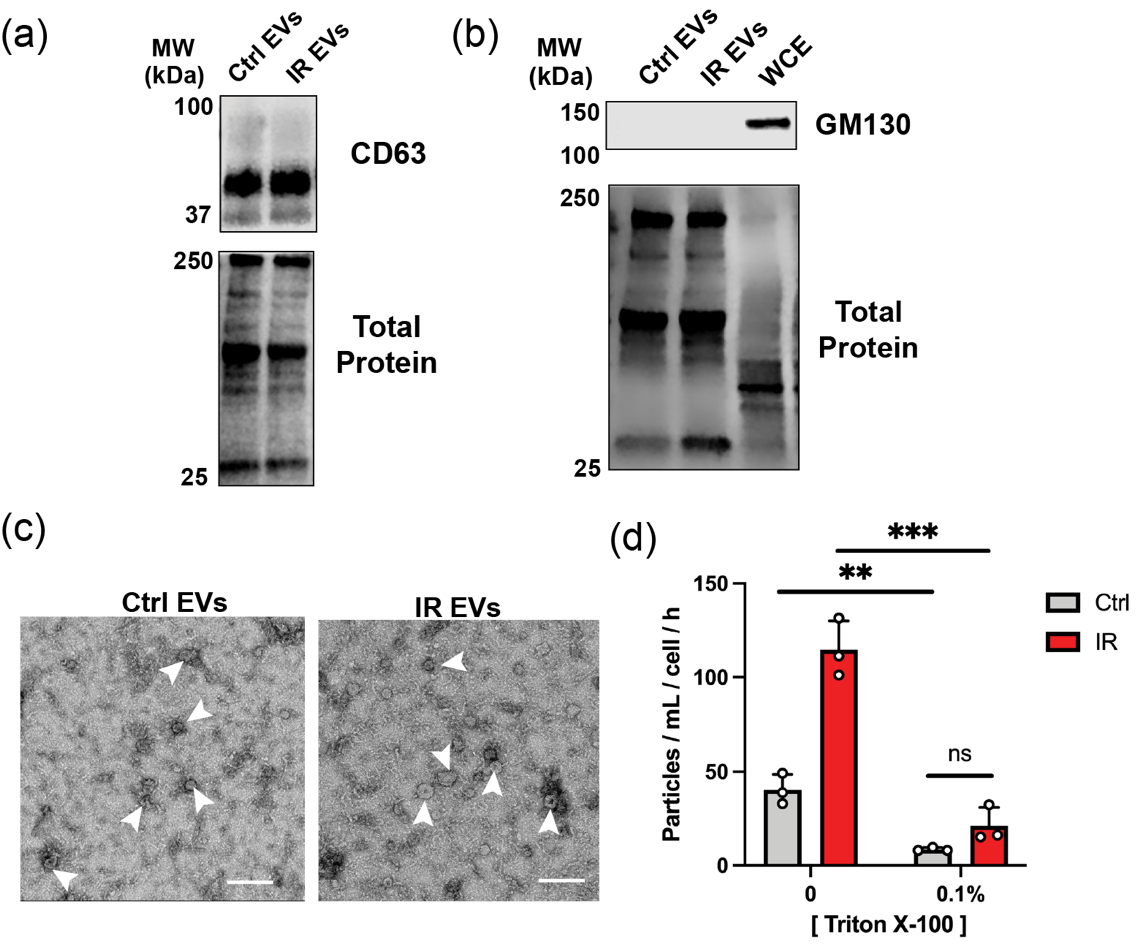

**Figure S1. Characterization of EV preparations derived from irradiated RMFs.** Representative western blot analysis for **(a)** CD63 and **(b)** GM130 expression in indicated protein extracts. **(c)** Representative transmission electron microscopy images of EVs derived from control (Ctrl; 0 Gy) and irradiated (IR; 10 Gy) RMFs. White arrowheads indicate small EVs. Scale bar represents 200 μm. **(d)** Particle concentration normalized to cell number and conditioning time for EV preparations from control and irradiated RMFs pre-treated with Triton X-100 at the indicated concentrations, determined via NTA. Statistical significance was determined via unpaired *t*-tests with Welch’s correction, with nonsignificant (ns) *p* > 0.5, ***p* < 0.01, and ****p* < 0.001. WCE = whole cell extract. Error bars show standard deviation.

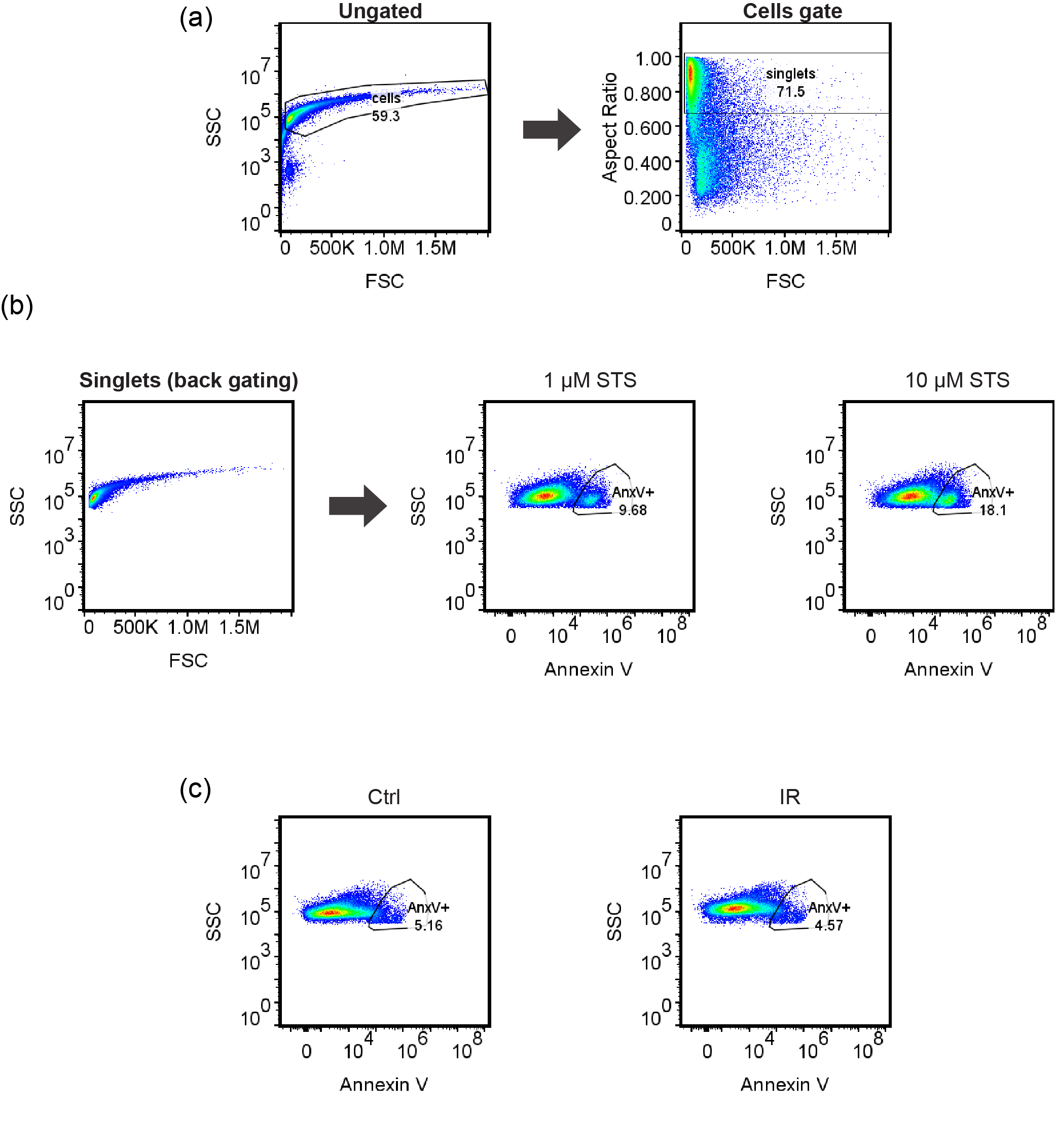

**Figure S2. Flow cytometry gating strategy for apoptosis analysis of irradiated RMFs. (a)** Gating strategy used to obtain single cell data (singlets). **(b)** Gating of singlets for the marker Annexin V to determine the population of apoptotic cells (AnxV+). RMFs treated with staurosporine (STS) at the indicated concentrations for 4 hours were used a positive control to set the AnxV+ gate. **(c)** AnxV+ gate applied to control (Ctrl; 0 Gy) and irradiated (IR; 10 Gy) singlets. Density plots show data from one representative experiment. FSC=forward scatter, SSC=side scatter.

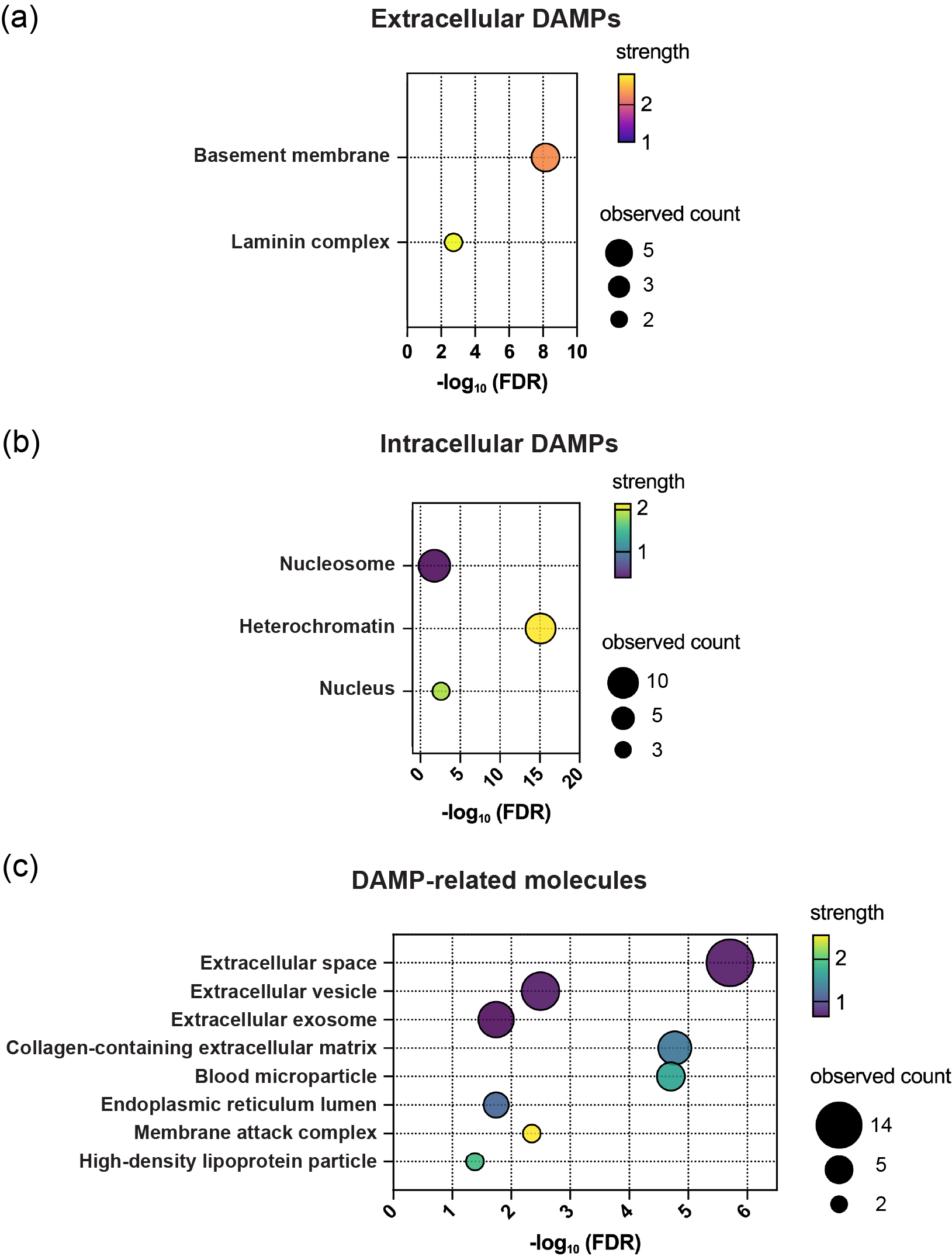

**Figure S3. GO cellular component analysis of indicated significant protein subsets identified via MS analysis of EVs derived from irradiated RMFs. (a)** Extracellular DAMPs. Proteins in network: AGRN, LAMB2, HSPG2, HMCN1, NTN4. **(b)** Intracellular DAMPs. Proteins in network: H2BC12, H2AZ1, H2AC1, H1-4, H2AX, H1-5, H1-2, H2BC26, H4, TAF15. **(c)** DAMP-related molecules. Proteins in network: C5, APOA1, C9, AMBP, AHSG, SERPINE2, ITIH2, FST, CD82, GPC1, FEN1, NRG1, TSKU, FKBP10, LOXL2, PLTP. X-axis quantification represents -log_10_ False Discovery Rate (FDR). Bubble size represents counts in network. Bubble color represents enrichment strength (log_10_ observed / expected). DAMPs = damage associated molecular patterns.

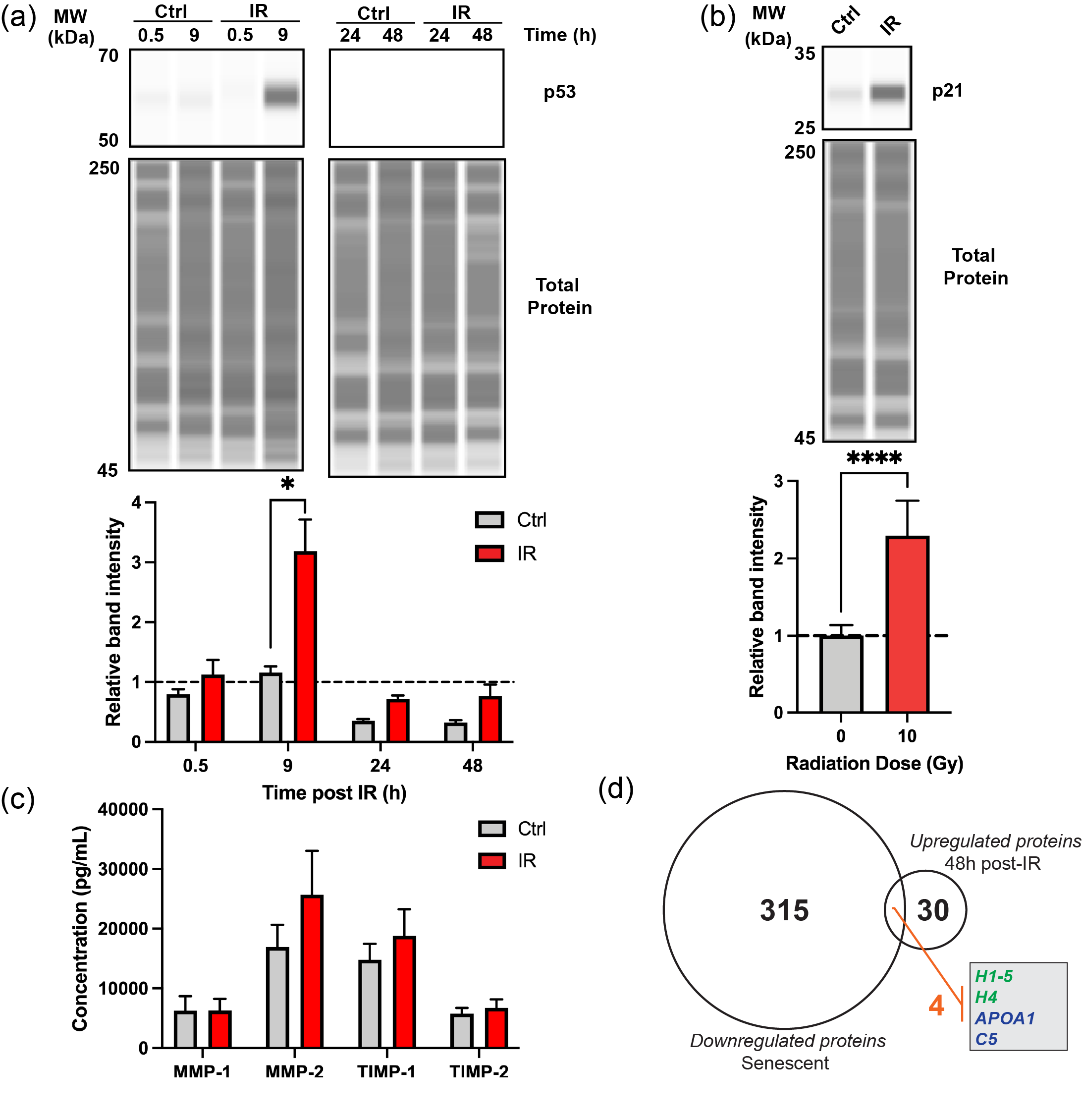

**Figure S4. Senescence characterization in irradiated RMFs. (a)** Representative western blot analysis and quantification of p53 expression in whole cell protein extracts from control (Ctrl; 0 Gy) and irradiated (IR; 10 Gy) RMFs at indicated time points post-IR. Total protein stain was utilized for normalization (N=3). **(b)** Representative western blot analysis and quantification of p21 expression in whole protein cell extracts from control and irradiated RMFs 48 hours post-IR. Total protein stain was utilized for normalization (N=3). **(c)** Concentration (pg/mL) of secreted MMP-1, MMP-2, TIMP-1, and TIMP-2 from control and irradiated RMF CM, determined via Luminex assay (N=3). **(d)** Venn diagram comparing proteins upregulated in EVs secreted by irradiated RMFs and those downregulated in senescent human fibroblast EVs (data from SASP atlas; [www.saspatlas.org](http://www.saspatlas.org); Basisty *et al.,* 2020). Statistical significance determined via unpaired *t*-tests with **p* < 0.05 and ****p* < 0.001. For statistical analysis of p21 expression, Welch’s correction was utilized. Error bars show standard deviation

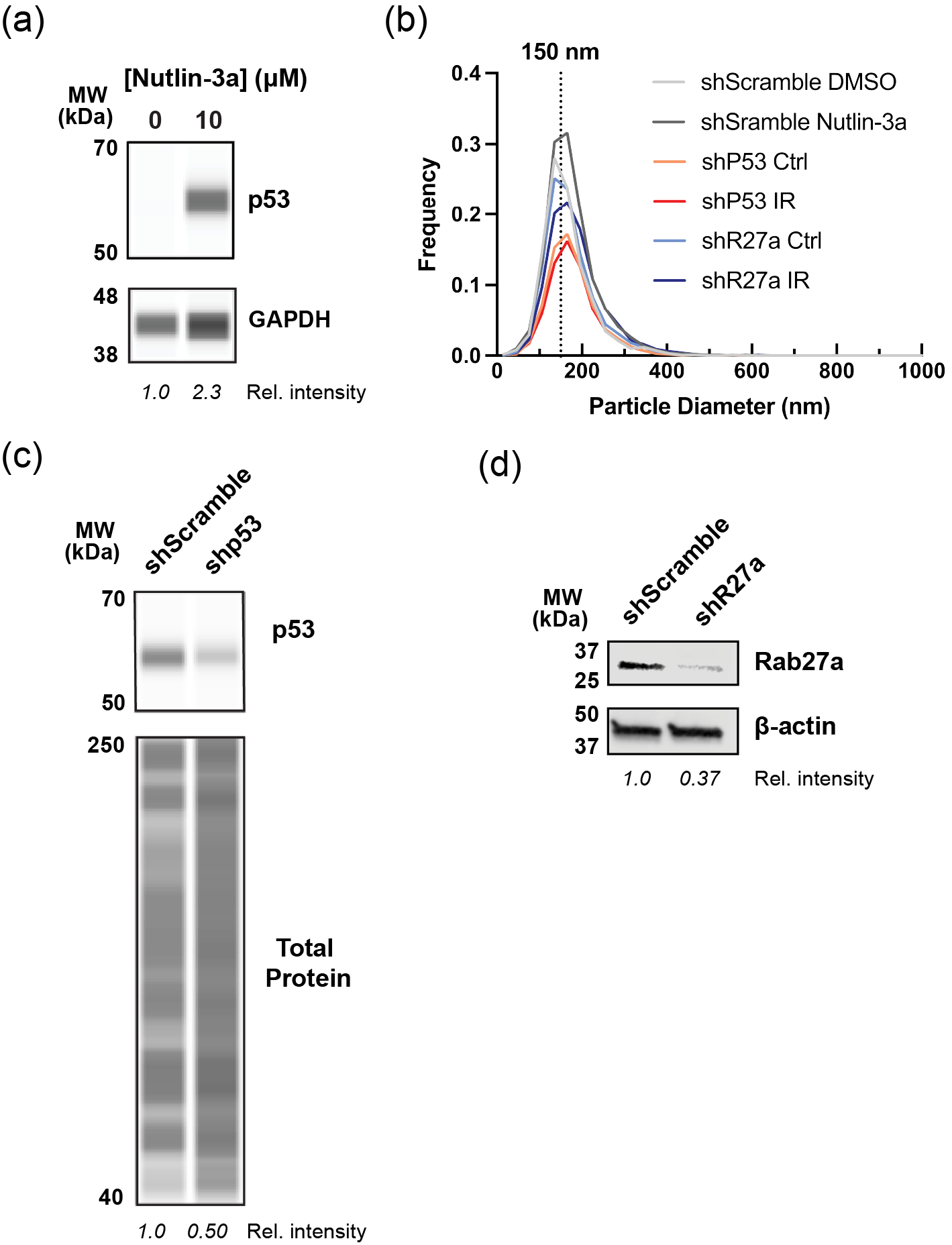

**Figure S5. Analysis of p53 activation, and knockdown of p53 and Rab27a in RMFs. (a)** Representative western blot analysis of p53 expression in whole cell extracts from RMFs treated with Nutlin-3a at the indicated concentrations. GAPDH expression was utilized as loading control. **(b)** Size distribution of EV preparations derived from indicated experimental conditions. Approximate peak diameter is indicated with dotted line. **(c)** Knockdown of p53 in RMFs. Representative western blot analysis of p53 expression in whole cell extracts from RMFs transformed with scramble control shRNA (shScramble) and p53 shRNA (shp53). Total protein stain was utilized as loading control. **(d)** Knockdown of Rab27a in RMFs. Representative western blot analysis of Rab27a expression in whole cell protein extracts from RMFs transformed with shScramble and Rab27a shRNA (shR27a). b-actin expression was utilized as loading control.

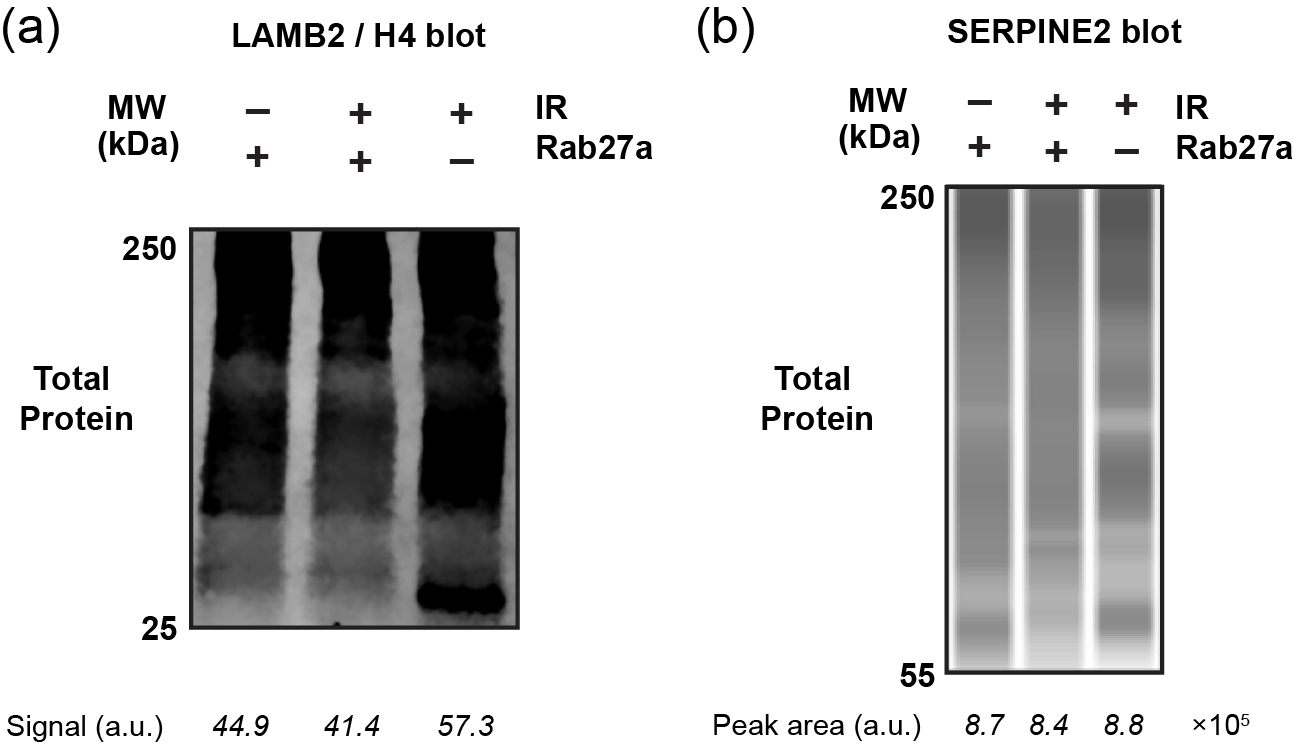

**Figure S6. Total protein stain from western blot analysis of selected DAMPs in EV-derived protein extracts**. Total protein stain corresponding to representative **(a)** LAMB2 / H4 and **(b)** SERPINE2 blots, depicted in Figure 4a. For each blot, equivalent protein amount of EV-derived extracts from indicated experimental conditions was loaded. Signal or peak area quantification of total protein stain is shown below each lane.

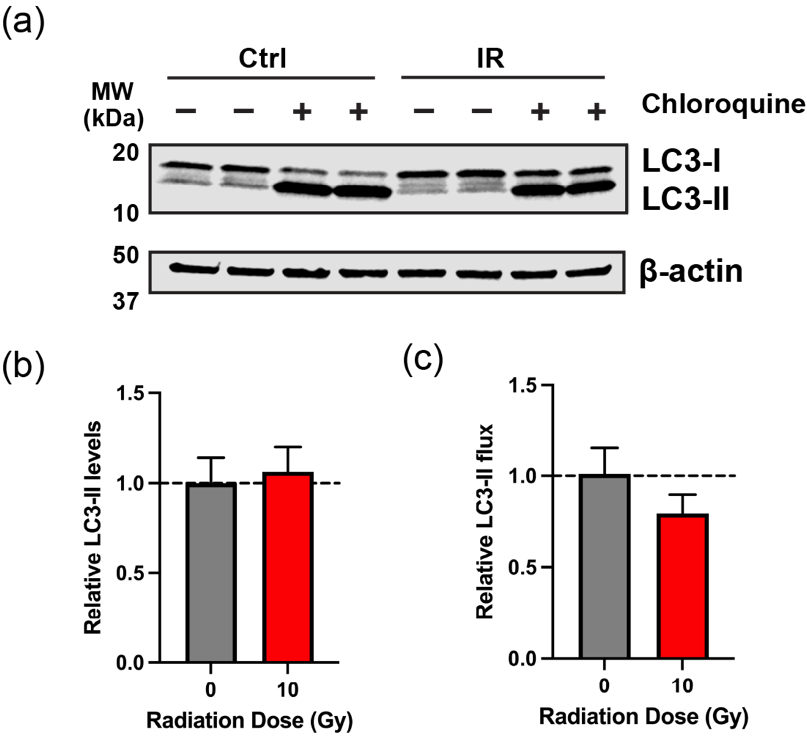

**Figure S7. Analysis of macroautophagy in irradiated RMFs. (a)** Representative western blot analysis of LC3 expression in whole cell extracts derived from control (Ctrl; 0 Gy) and irradiated (IR; 10 Gy) RMFs. β-actin was utilized as loading control. Cells were supplemented with chloroquine for 4 hours where indicated (+) to inhibit autophagy. **(b)** Quantification of steady-state LC3-II values 48 hours post-IR. Data are normalized to b-actin expression and represent normalized LC3-II expression relative to the control average (dotted line). **(c)** Quantification of LC3-II values with inhibition of autophagy. Flux was determined by ratio of normalized LC3-II expression of cells supplemented with chloroquine to the normalized LC3-II value at steady state. Flux values are relative to control average (dotted line). Error bars show standard deviation.

**SUPPLEMENTAL TABLES**

**Table S1. List of protein markers analyzed in the 15-plex Luminex assay.**

| Marker | Description |
| --- | --- |
| GM-CSF | Granulocyte-macrophage colony-stimulating factor |
| IFN-γ | Interferon gamma |
| GM-CSF | Granulocyte-macrophage colony-stimulating factor |
| IFNγ | Interferon gamma |
| IL-1β | Interleukin-1 beta |
| IL-1Ra | Interleukin-1 receptor antagonist |
| IL-2 | Interleukin-2 |
| IL-4 | Interleukin-4 |
| IL-5 | Interleukin-5 |
| IL-6 | Interleukin-6 |
| IL-8 | Interleukin-8 (CXCL8) |
| IL-10 | Interleukin-10 |
| IL-12p40 | Interleukin-12 subunit p40 |
| IL-12p70 | Interleukin-12 subunit p70 (active heterodimer) |
| IL-13 | Interleukin-13 |
| MCP-1 | Monocyte chemoattractant protein-1 (CCL2) |
| TNF-α | Tumor necrosis factor alpha |

**Table S2. List of protein markers analyzed in the 13-plex Luminex assay.**

| Marker | Description |
| --- | --- |
| MMP-1 | Matrix metalloproteinase 1 |
| MMP-2 | Matrix metalloproteinase 2 |
| MMP-3 | Matrix metalloproteinase 3 |
| MMP-7 | Matrix metalloproteinase 7 |
| MMP-8 | Matrix metalloproteinase 8 |
| MMP-9 | Matrix metalloproteinase 9 |
| MMP-10 | Matrix metalloproteinase 10 |
| MMP-12 | Matrix metalloproteinase 12 |
| MMP-13 | Matrix metalloproteinase 13 |
| TIMP-1 | Tissue inhibitors of metalloproteinases 1 |
| TIMP-2 | Tissue inhibitors of metalloproteinases 2 |
| TIMP-3 | Tissue inhibitors of metalloproteinases 3 |
| TIMP-4 | Tissue inhibitors of metalloproteinases 4 |

**Table S3. List of proteins quantified through tandem mass tag-based MS analysis of EVs derived from irradiated RMFs.**

Supplied as an Excel file.

**Table S4. Gene ontology of significant upregulated proteins determined through MS analysis of irradiated RMF EVs.**

| Term ID | Category | Term description | Observed gene count | Background gene count | Strength | False discovery rate | Matching proteins in network |
| --- | --- | --- | --- | --- | --- | --- | --- |
| GO:0030527 | Function | Structural constituent of chromatin | 9 | 101 | 1.71 | 8.58E-10 | H4C6, H2AZ1, H1-4, H1-5, H1-2, H2BC12, H2AC13, H2AX, H2BU1 |
| GO:0005198 | Function | Structural molecule activity | 13 | 776 | 0.99 | 6.36E-07 | H4C6, HMCN1, H2AZ1, LAMB2, H1-4, H1-5, H1-2, H2BC12, H2AC13, HSPG2, AGRN, H2AX, H2BU1 |
| GO:0031492 | Function | Nucleosomal DNA binding | 4 | 23 | 2 | 0.00021 | H2AZ1, H1-4, H1-5, H1-2 |
| GO:0004866 | Function | Endopeptidase inhibitor activity | 6 | 177 | 1.29 | 0.00078 | C5, SERPINA10, AMBP, AHSG, ITIH2, SERPINE2 |
| GO:0044877 | Function | Protein-containing complex binding | 11 | 1261 | 0.7 | 0.003 | APOA1, FST, AMBP, NRG1, H2AZ1, LAMB2, H1-4, H1-5, H1-2, HSPG2, PLTP |
| GO:0043236 | Function | Laminin binding | 3 | 28 | 1.79 | 0.01 | GPC1, NTN4, AGRN |
| GO:0004867 | Function | Serine-type endopeptidase inhibitor activity | 4 | 98 | 1.37 | 0.0124 | SERPINA10, AMBP, ITIH2, SERPINE2 |
| GO:0046982 | Function | Protein heterodimerization activity | 6 | 368 | 0.98 | 0.0145 | H4C6, H2AZ1, H2BC12, H2AC13, H2AX, H2BU1 |
| GO:0005539 | Function | Glycosaminoglycan binding | 5 | 245 | 1.07 | 0.0212 | SERPINA10, AMBP, ITIH2, AGRN, SERPINE2 |
| GO:1901681 | Function | Sulfur compound binding | 5 | 272 | 1.03 | 0.0323 | FST, SERPINA10, AMBP, AGRN, SERPINE2 |
| GO:0006334 | Process | Nucleosome assembly | 7 | 124 | 1.51 | 3.51E-05 | H4C6, H1-4, H1-5, H1-2, H2BC12, H2AX, H2BU1 |
| GO:0051346 | Process | Negative regulation of hydrolase activity | 7 | 354 | 1.06 | 0.0051 | C5, APOA1, SERPINA10, AMBP, AHSG, ITIH2, SERPINE2 |
| GO:0010605 | Process | Negative regulation of macromolecule metabolic process | 16 | 2760 | 0.53 | 0.0067 | C5, APOA1, FST, SERPINA10, AMBP, AHSG, NRG1, H1-4, H1-5, H1-2, ITIH2, HSPG2, LOXL2, SERPINE2, TSKU, TAF15 |
| GO:0043933 | Process | Protein-containing complex organization | 12 | 1465 | 0.68 | 0.0067 | APOA1, H4C6, C9, NRG1, H1-4, H1-5, H1-2, H2BC12, SV2A, PLTP, H2AX, H2BU1 |
| GO:0022607 | Process | Cellular component assembly | 15 | 2467 | 0.55 | 0.0069 | APOA1, H4C6, C9, GPC1, NRG1, LAMB2, H1-4, FKBP10, H1-5, H1-2, NTN4, H2BC12, SV2A, H2AX, H2BU1 |
| GO:0006325 | Process | Chromatin organization | 8 | 612 | 0.88 | 0.0087 | H4C6, H1-4, H1-5, H1-2, H2BC12, LOXL2, H2AX, H2BU1 |
| GO:0016043 | Process | Cellular component organization | 22 | 5436 | 0.37 | 0.0087 | APOA1, H4C6, C9, GPC1, HMCN1, NRG1, FEN1, LAMB2, H1-4, FKBP10, H1-5, H1-2, NTN4, H2BC12, SV2A, AGRN, LOXL2, SERPINE2, PLTP, H2AX, TSKU, H2BU1 |
| GO:0065003 | Process | Protein-containing complex assembly | 11 | 1303 | 0.69 | 0.0087 | APOA1, H4C6, C9, NRG1, H1-4, H1-5, H1-2, H2BC12, SV2A, H2AX, H2BU1 |
| GO:0051172 | Process | Negative regulation of nitrogen compound metabolic process | 14 | 2403 | 0.53 | 0.0172 | C5, FST, SERPINA10, AMBP, AHSG, NRG1, H1-4, H1-5, H1-2, ITIH2, HSPG2, LOXL2, SERPINE2, TAF15 |
| GO:0071711 | Process | Basement membrane organization | 3 | 30 | 1.76 | 0.0191 | HMCN1, LAMB2, NTN4 |
| GO:0051276 | Process | Chromosome organization | 9 | 968 | 0.73 | 0.021 | H4C6, FEN1, H1-4, H1-5, H1-2, H2BC12, LOXL2, H2AX, H2BU1 |
| GO:0050793 | Process | Regulation of developmental process | 14 | 2492 | 0.51 | 0.0215 | C5, APOA1, H4C6, FST, GPC1, AHSG, NRG1, LAMB2, NTN4, HSPG2, AGRN, LOXL2, SERPINE2, TSKU |
| GO:0085029 | Process | Extracellular matrix assembly | 3 | 35 | 1.7 | 0.0247 | LAMB2, FKBP10, NTN4 |
| GO:0014037 | Process | Schwann cell differentiation | 3 | 39 | 1.65 | 0.0321 | GPC1, NRG1, LAMB2 |
| GO:0030261 | Process | Chromosome condensation | 3 | 42 | 1.62 | 0.0381 | H1-4, H1-5, H1-2 |
| GO:0045910 | Process | Negative regulation of DNA recombination | 3 | 44 | 1.6 | 0.0418 | H1-4, H1-5, H1-2 |
| GO:0005615 | Component | Extracellular space | 25 | 3247 | 0.65 | 6.53E-10 | C5, CD82, APOA1, H4C6, FST, SERPINA10, C9, GPC1, AMBP, HMCN1, AHSG, NRG1, H2AZ1, LAMB2, H2BC12, ITIH2, H2AC13, HSPG2, AGRN, LOXL2, MOGS, SERPINE2, PLTP, H2AX, TSKU |
| GO:0000786 | Component | Nucleosome | 9 | 133 | 1.59 | 1.85E-09 | H4C6, H2AZ1, H1-4, H1-5, H1-2, H2BC12, H2AC13, H2AX, H2BU1 |
| GO:0062023 | Component | Collagen-containing extracellular matrix | 12 | 407 | 1.23 | 1.85E-09 | APOA1, GPC1, AMBP, HMCN1, AHSG, LAMB2, NTN4, ITIH2, HSPG2, AGRN, LOXL2, SERPINE2 |
| GO:0005576 | Component | Extracellular region | 26 | 4175 | 0.56 | 4.81E-09 | C5, CD82, APOA1, H4C6, FST, SERPINA10, C9, GPC1, AMBP, HMCN1, AHSG, NRG1, H2AZ1, LAMB2, NTN4, H2BC12, ITIH2, H2AC13, HSPG2, AGRN, LOXL2, MOGS, SERPINE2, PLTP, H2AX, TSKU |
| GO:1903561 | Component | Extracellular vesicle | 19 | 2120 | 0.72 | 4.08E-08 | C5, CD82, APOA1, H4C6, SERPINA10, C9, GPC1, AMBP, HMCN1, AHSG, H2AZ1, LAMB2, ITIH2, H2AC13, HSPG2, AGRN, MOGS, SERPINE2, H2AX |
| GO:0070062 | Component | Extracellular exosome | 18 | 2096 | 0.7 | 2.24E-07 | C5, CD82, APOA1, H4C6, SERPINA10, C9, GPC1, AMBP, HMCN1, AHSG, H2AZ1, LAMB2, ITIH2, H2AC13, HSPG2, AGRN, MOGS, H2AX |
| GO:0005604 | Component | Basement membrane | 6 | 98 | 1.55 | 3.48E-06 | HMCN1, LAMB2, NTN4, HSPG2, AGRN, LOXL2 |
| GO:0031982 | Component | Vesicle | 20 | 3957 | 0.47 | 0.00012 | C5, CD82, APOA1, H4C6, SERPINA10, C9, GPC1, AMBP, HMCN1, AHSG, H2AZ1, LAMB2, ITIH2, H2AC13, SV2A, HSPG2, AGRN, MOGS, SERPINE2, H2AX |
| GO:0072562 | Component | Blood microparticle | 5 | 118 | 1.39 | 0.00028 | APOA1, C9, AMBP, AHSG, ITIH2 |
| GO:0031594 | Component | Neuromuscular junction | 4 | 73 | 1.5 | 0.0012 | NRG1, LAMB2, SV2A, SERPINE2 |
| GO:0005788 | Component | Endoplasmic reticulum lumen | 6 | 312 | 1.05 | 0.0018 | APOA1, SERPINA10, AHSG, LAMB2, FKBP10, ITIH2 |
| GO:0000785 | Component | Chromatin | 10 | 1285 | 0.65 | 0.005 | H4C6, H2AZ1, H1-4, H1-5, H1-2, H2BC12, H2AC13, LOXL2, H2AX, H2BU1 |
| GO:0005579 | Component | Membrane attack complex | 2 | 7 | 2.22 | 0.0111 | C5, C9 |
| GO:0005694 | Component | Chromosome | 11 | 1850 | 0.54 | 0.019 | H4C6, H2AZ1, FEN1, H1-4, H1-5, H1-2, H2BC12, H2AC13, LOXL2, H2AX, H2BU1 |
| GO:0043227 | Component | Membrane-bounded organelle | 32 | 13188 | 0.15 | 0.019 | C5, CD82, APOA1, H4C6, FST, SERPINA10, C9, GPC1, AMBP, HMCN1, AHSG, NRG1, H2AZ1, FEN1, LAMB2, H1-4, FKBP10, H1-5, H1-2, H2BC12, ITIH2, H2AC13, SV2A, HSPG2, AGRN, LOXL2, MOGS, SERPINE2, PLTP, H2AX, TAF15, H2BU1 |
| GO:0043256 | Component | Laminin complex | 2 | 12 | 1.98 | 0.0241 | LAMB2, NTN4 |
| GO:0000792 | Component | Heterochromatin | 3 | 77 | 1.35 | 0.0312 | H2AZ1, H1-4, H1-5 |

**Table S5. List of Ras-associated binding (Rab) proteins identified via MS analysis of RMF EVs.**

| Accession | Protein name |
| --- | --- |
| Q9H0U4 | Rab1b |
| P62820 | Rab1a |
| P51153 | Rab13 |
| Q92930 | Rab8b |
| P51159 | Rab27a |
| P61026 | Rab10 |
| P20337 | Rab3b |
| P51148 | Rab5c |
| Q9NP72 | Rab18 |
| Q15286 | Rab35 |
| Q15907 | Rab11b |
| P51151 | Rab9a |
| Q9BZG1 | Rab34 |
| P61019 | Rab2a |
| Q9UL25 | Rab21 |
| Q13637 | Rab32 |
| P20339 | Rab5a |
| P61106 | Rab14 |
| Q9ULC3 | Rab23 |
| P51149 | Rab7a |
